# p*K*_a_ measurements for the SAMPL6 prediction challenge for a set of kinase inhibitor-like fragments

**DOI:** 10.1101/368787

**Authors:** Mehtap Işik, Dorothy Levorse, Ariën S. Rustenburg, Ikenna E. Ndukwe, Heather Wang, Xiao Wang, Mikhail Reibarkh, Gary E. Martin, Alexey A. Makarov, David L. Mobley, Timothy Rhodes, John D. Chodera

**Affiliations:** Computational and Systems Biology Program, Sloan Kettering Institute, Memorial Sloan Kettering Cancer Center, New York, NY 10065, United States; Tri-Institutional PhD Program in Chemical Biology, Weill Cornell Graduate School of Medical Sciences, Cornell University, New York, NY 10065, United States; Pharmaceutical Sciences, MRL, Merck & Co., Inc., 126 East Lincoln Avenue, Rahway, New Jersey 07065, United States; Graduate Program in Physiology, Biophysics, and Systems Biology, Weill Cornell Medical College, New York, NY 10065, United States; Process and Analytical Research and Development, Merck & Co., Inc., Rahway, NJ 07065, United States; Analytical Research & Development, MRL, Merck & Co., Inc., MRL, 126 East Lincoln Avenue, Rahway, New Jersey 07065, United States; Department of Pharmaceutical Sciences and Department of Chemistry, University of California, Irvine, Irvine, California 92697, United States

**Author notes:** (TR); (JDC).

**Keywords:** acid dissociation constants, spectrophotometric p*K*_a_ measurement, blind prediction challenge, SAMPL, macroscopic p*K*_a_, microscopic p*K*_a_, macroscopic protonation state, microscopic protonation state

## Abstract

Determining the net charge and protonation states populated by a small molecule in an environment of interest or the cost of altering those protonation states upon transfer to another environment is a prerequisite for predicting its physicochemical and pharmaceutical properties. The environment of interest can be aqueous, an organic solvent, a protein binding site, or a lipid bilayer. Predicting the protonation state of a small molecule is essential to predicting its interactions with biological macromolecules using computational models. Incorrectly modeling the dominant protonation state, shifts in dominant protonation state, or the population of significant mixtures of protonation states can lead to large modeling errors that degrade the accuracy of physical modeling. Low accuracy hinders the use of physical modeling approaches for molecular design. For small molecules, the acid dissociation constant (p*K*_a_) is the primary quantity needed to determine the ionic states populated by a molecule in an aqueous solution at a given pH. As a part of SAMPL6 community challenge, we organized a blind p*K*_a_ prediction component to assess the accuracy with which contemporary p*K*_a_ prediction methods can predict this quantity, with the ultimate aim of assessing the expected impact on modeling errors this would induce. While a multitude of approaches for predicting p*K*_a_ values currently exist, predicting the p*K*_a_s of drug-like molecules can be difficult due to challenging properties such as multiple titratable sites, heterocycles, and tautomerization. For this challenge, we focused on set of 24 small molecules selected to resemble selective kinase inhibitors—an important class of therapeutics replete with titratable moieties. Using a Sirius T3 instrument that performs automated acid-base titrations, we used UV absorbance-based p*K*_a_ measurements to construct a high-quality experimental reference dataset of macroscopic p*K*_a_s for the evaluation of computational p*K*_a_ prediction methodologies that was utilized in the SAMPL6 p*K*_a_ challenge. For several compounds in which the microscopic protonation states associated with macroscopic p*K*_a_s were ambiguous, we performed follow-up NMR experiments to disambiguate the microstates involved in the transition. This dataset provides a useful standard benchmark dataset for the evaluation of p*K*_a_ prediction methodologies on kinase inhibitor-like compounds.

**Abbreviations:**

SAMPL: Statistical Assessment of the Modeling of Proteins and Ligands
p*K*_a_: -log_10_ acid dissociation equilibrium constant
p_s_*K*_a_: -log_10_ apparent acid dissociation equilibrium constant in cosolvent
DMSO: Dimethyl sulfoxide
ISA: lonic-strength adjusted
SEM: Standard error of the mean
TFA: Target factor analysis
LC-MS: Liquid chromatography - mass spectrometry
NMR: Nuclear magnetic resonance spectroscopy
HMBC: Heteronuclear Multiple-Bond Correlation
TFA-*d*: deutero-trifluoroacetic acid

## Introduction

SAMPL (Statistical Assessment of the Modeling of Proteins and Ligands) is a recurring series of blind prediction challenges for the computational chemistry community [1, 2]. Through these challenges, SAMPL aims to evaluate and advance computational tools for rational drug design. SAMPL has driven progress in a number of areas over seven previous rounds of challenge cycles [3–7, 7–15] by focusing the community on specific phenomena relevant to drug discovery poorly predicted by current models, isolating that phenomenon from other confounding factors in well-designed test systems, evaluating tools prospectively, enabling data sharing to learn from failures, and releasing the resulting high-quality datasets into the community as benchmark sets.

As a stepping stone to enabling the accurate prediction of protein-ligand binding affinities, SAMPL has focused on evaluating how well physical and empirical modeling methodologies can predict various physicochemical properties relevant to binding and drug discovery, such as hydration free energies (which model aspects of desolvation in isolation), distribution coefficients (which model transfer from relatively homogeneous aqueous to nonpolar environments), and host-guest binding affinities (which model high-affinity association without the complication of slow protein dynamics). These physicochemical property prediction challenges—in addition to assessing the predictive accuracy of quantities that are useful in various stages of drug discovery in their own right—have been helpful in pinpointing deficiencies in computational models that can lead to substantial errors in affinity predictions.

### Neglect of protonation state effects can lead to large modeling errors

As part of the SAMPL5 challenge series, a new cyclohexane-water distribution constant (log *D*) prediction challenge was introduced, where participants predicted the transfer free energy of small drug-like molecules between an aqueous buffer phase at pH 7.4 and a nonaqueous cyclohexane phase [16, 17]. While octanol-water distribution coefficient measurements are more common, cyclohexane was selected for the simplicity of its liquid phase and relative dryness compared to wet octanol phases. While the expectation was that this challenge would be relatively straightforward given the lack of complexity of cyclohexane phases, analysis of participant performance revealed that multiple factors contributed to significant prediction failures: poor conformational sampling of flexible solute molecules, misprediction of relevant protonation and tautomeric states (or failure to accommodate shifts in their populations), and force field inaccuracies resulting in bias towards the cyclohexane phase. While these findings justified the benefit of future iterations of blind distribution or partition coefficient challenges, the most surprising observation from this initial log *D* challenge was that participants almost uniformly neglected to accurately model protonation state effects, and that neglect of these effects led to surprisingly large errors in transfer free energies [16–18]. Careful quantum chemical assessments of the magnitude of these protonation state effects found that their neglect could introduce errors up to 6–8 kcal/mol for some compounds [18]. This effect stems from the need to account for the free energy difference between the major ionization state in cyclohexane (most likely neutral state) and in water phase (which could be neutral or charged).

To isolate these surprisingly large protonation state modeling errors from difficulties related to lipophilicity (log *P* and log *D*) prediction methods, we decided to organize a set of staged physicochemical property challenges using a consistent set of molecules that resemble small molecule kinase inhibitors—an important drug class replete with multiple titratable moieties. This series of challenges will first evaluate the ability of current-generation modeling tools to predict acid dissociation constants (p*K*_a_). It will be followed by a partition/distribution coefficient challenge to evaluate the ability to incorporate experimentally-provided p*K*_a_ values into prediction of distribution coefficients to ensure methodologies can correctly incorporate protonation state effects into their predictions. A third challenge stage will follow: a new blinded partition/distribution coefficient challenge where participants must predict p*K*_a_ values on their own. At the conclusion of this series of challenges, we will ensure that modern physical and empirical modeling methods have eliminated this large source of spurious errors from modeling both simple and complex phenomena.

This article reports on the experiments for the first stage of this series of challenges: SAMPL6 p*K*_a_ prediction challenge. The selection of a small molecule set and collection of experimental p*K*_a_ data are described in detail.

### Conceptualization of a blind p*K*_a_ challenge

This is the first time a blind p*K*_a_ prediction challenge has been fielded as part of SAMPL. In this first iteration of the challenge, we aimed to assess the performance of current p*K*_a_ prediction methods and isolate potential causes of inaccurate p*K*_a_ estimates.

The prediction of p*K*_a_ values for drug-like molecules can be complicated by several effects: the presence of multiple (potentially coupled) titratable sites, the presence of heterocycles, tautomerization, the conformational flexibility of large molecules, and ability of intramolecular hydrogen bonds to form. We decided to focus on the chemical space of small molecule kinase inhibitors in the first iteration of p*K*_a_ prediction challenge. A total of 24 small organic molecules (17 drug-fragment-like and 7 drug-like) were selected for their similarity to known small molecule kinase inhibitors, while also considering properties predicted to affect the experimental tractability of p*K*_a_ and log *P* measurements such as solubility and predicted p*K*_a_s. Macroscopic p*K*_a_ values were collected experimentally with UV-absorbance spectroscopy-based p*K*_a_ measurements using a Sirius T3 instrument, which automates the sample handling, titration, and spectroscopic measurements to allow high-quality p*K*_a_ determination. The Sirius T3 is equipped with an autosampler which allowed us to run 8–10 measurements per day. Experimental data were kept blinded for three months (25 Oct 2017 through 23 Jan 2018) to allow participants in the SAMPL6 p*K*_a_ challenge to submit truly blinded computational predictions. Eleven research groups participated in this challenge, providing a total of 93 prediction submission sets that cover a large variety of contemporary p*K*_a_ prediction methods.

### Our selected experimental approach determines macroscopic p*K*_a_ values

Whenever experimental p*K*_a_ measurements are used for evaluating p*K*_a_ predictions, it is important to differentiate between microscopic and macroscopic p*K*_a_ values. In molecules containing multiple titratable moieties, the protonation state of one group can affect the proton dissociation propensity of another functional group. In such cases, the **microscopic p*K*_*a*_** (group p*K*_a_) refers to the p*K*_a_ of deprotonation of a single titratable group while all the other titratable and tautomerizable functional groups of the same molecule are held fixed. Different protonation states and tautomer combinations constitute different microstates. The **macroscopic p*K*_a_** (molecular p*K*_a_) defines the acid dissociation constant related to the observable loss of a proton from a molecule regardless of which functional group the proton is dissociating from, so it doesn’t necessarily convey structural information.

Whether a measured p*K*_a_ is microscopic or macroscopic depends on the experimental method used (Figure 2). For a molecule with only one titratable proton, the microscopic p*K*_a_ is equal to the macroscopic p*K*_a_. For a molecule with multiple titratable groups, however, throughout a titration from acidic to basic pH, the deprotonation of some functional groups can take place almost simultaneously. For these multiprotic molecules, the experimentally-measured macroscopic p*K*_a_ will include contributions from multiple microscopic p*K*_a_s with similar values (i.e., acid dissociation of multiple microstates). Cysteine provides an example of this behavior with its two macroscopic p*K*_a_s observable by spectrophotometric or potentiometric p*K*_a_ measurement experiments [19, 20].

**Figure 1.**
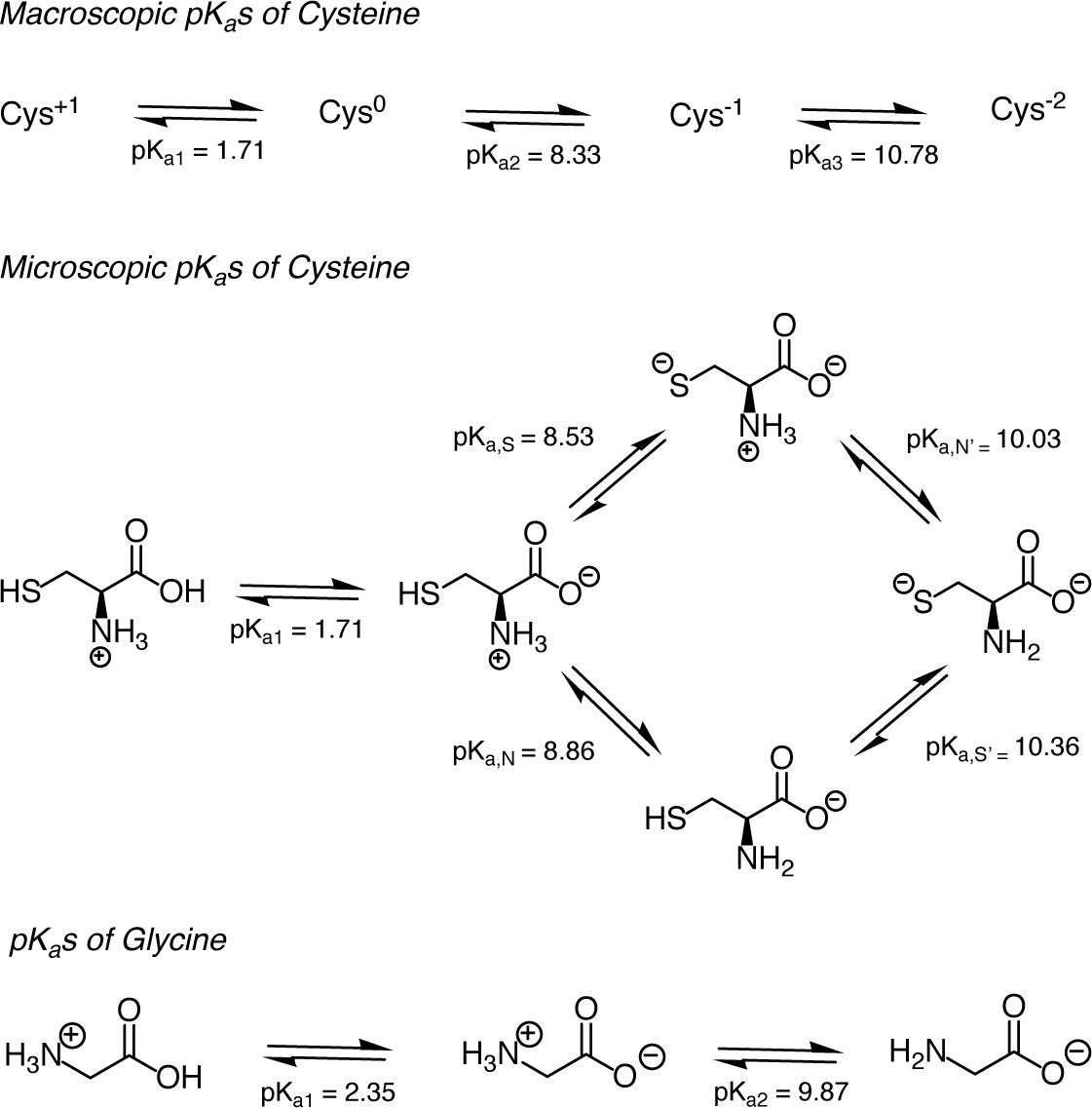
Assignment of cysteine and glycine p*K*_a_ values. ***pK***_***a*1**_, ***pK***_***a*2**_, and ***pK***_***a*3**_ are macroscopic acid dissociation constants for cysteine and glycine [24]. When p*K*_a_ values of a polyprotic molecule are very different, such as in the case of glycine, it is possible to assign the p*K*_a_s to individual groups since the dissociation of protons is stepwise [19]. However, stepwise dissociation cannot be assumed for cysteine, because ***pK***_***a*2**_ and ***pK***_***a*3**_ are very close in value. Four underlying microscopic p*K*_a_s (***pK***_***a,S***_, ***pK***_***a,N***_, ***pK***_***a,S***_, and ***pK***_***a,N′***_) for cysteine were measured using UV spectra analysis of cysteine and derivatives [25]. Notice that the proximity of ***pK***_***a,S***_ and ***pK***_***a,N***_ values indicates similar probability of proton dissociation from these groups. This figure is adopted from [19].

**Figure 2.**
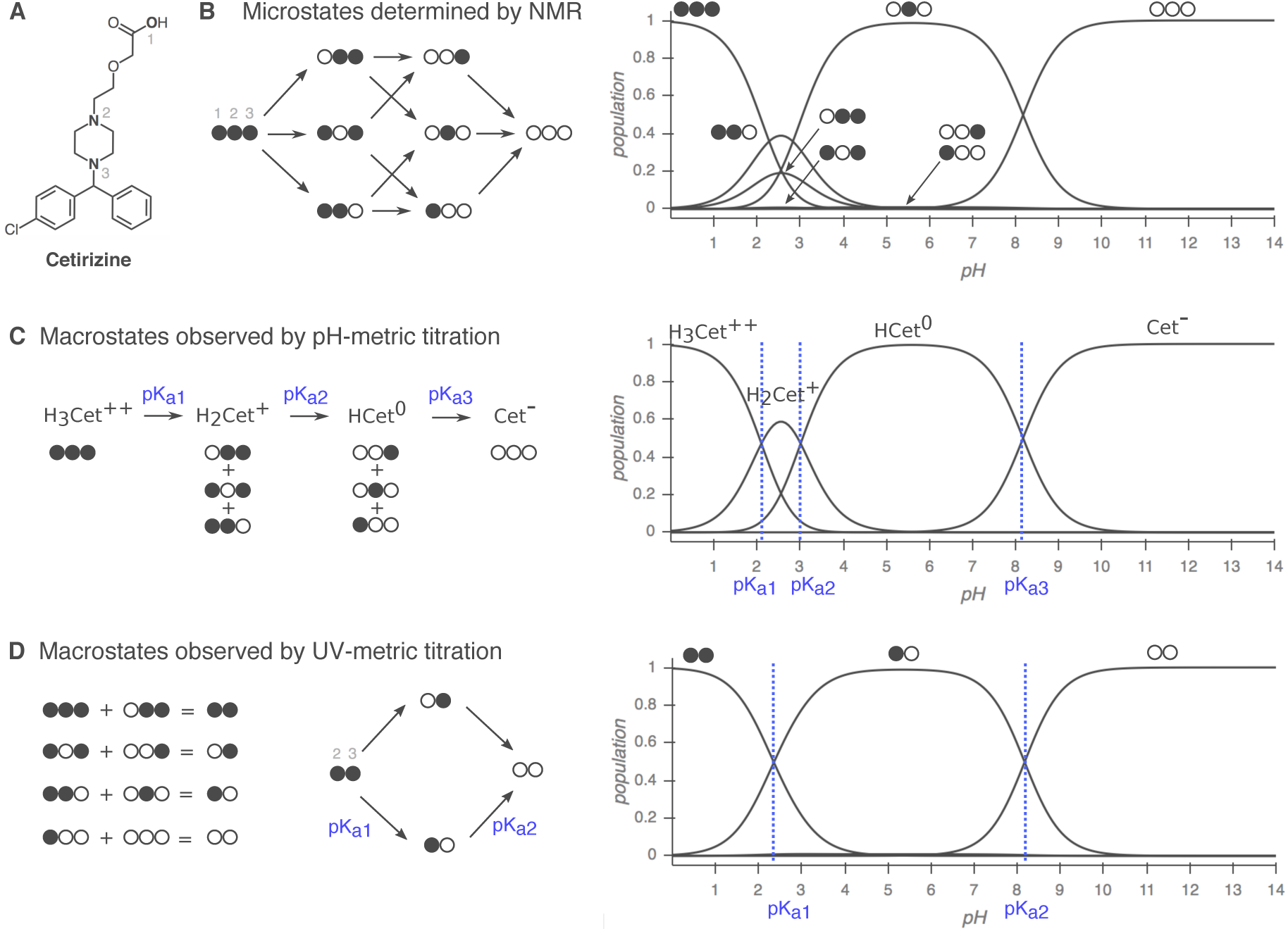
Comparison of macroscopic and microscopic p*K*_a_ measurement methods. Filled circles represent protonated sites and empty circles represent deprotonated sites with the order of carboxylic acid (1), piperazine nitrogen (2), and piperazine nitrogen (3). Protonation state populations shown for pH-metric and UV-metric p*K*_a_ measurement methods are simulations, calculated using NMR-based microscopic p*K*_a_ values. (**A**) Cetirizine has *n* =3 titratable sites, shown in bold. (**B**) *Left*: The 8 microstates (2^*n*^) and 12 microscopic p*K*_a_s (*n*2^*n*-1^) of cetirizine. *Right:* Relative population of microspecies with respect to pH. Potentially all microstates can be resolved via NMR. (**C**) Simulated pH-metric (potentiometric) titration and macroscopic populations. For a polyprotic molecule, only macroscopic p*K*_a_s can be measured with pH-metric titration. Microstates with different total charge (related to the number of protons) can be resolved, but microstates with the same total charge are observed as one macroscopic population. (**D**) Simulated microscopic populations for UV-metric (spectrophotometric) titration of cetirizine. Since only protonation of the titration sites within four heavy atoms of the UV-chromophore is likely to cause an observable change in the UV-absorbance spectra, microstates that only differ by protonation of the distal carboxylic acid cannot be differentiated. Moreover, populations that overlap may or may not be resolvable depending on how much their absorbance spectra in the UV region differ. Both UV-metric and pH-metric p*K*_a_ determination methods measure macroscopic p*K*_a_s for polyprotic molecules, which cannot easily be assigned to individual titration sites and underlying microstate populations in the absence of other experimental evidence that provides structural resolution, such as NMR. Note that macroscopic populations observed in these two methods are composed of different combinations of microstates depending on the principles of measurement technique. Here, the illustrative diagram style was adopted from [26], and NMR-determined microscopic p*K*_a_s for cetirizine were taken from [27].

While four microscopic p*K*_a_s can be defined for cysteine, experimentally observed p*K*_a_ values cannot be assigned to individual functional groups directly (Figure 1, *top*). More advanced techniques capable of resolving individual protonation sites—such as NMR [21], Raman spectroscopy [22, 23], and the analysis of p*K*_a_s in molecular fragments or derivatives—are required to unambiguously assign the site of protonation state changes. On the other hand, when there is a large difference between microscopic p*K*_a_s in a multiprotic molecule, the proton dissociations won’t overlap and macroscopic p*K*_a_s observed by experiments can be assigned to individual titratable groups. The p*K*_a_ values of glycine provide a good example of this scenario (Figure 1, *bottom*) [19, 20, 22]. We recommend the short review on the assignment of p*K*_a_ values authored by Ivan G. Darvey [20] for a good introduction to the concepts of macroscopic vs microscopic p*K*_a_ values.

The most common methods for measuring small molecule p*K*_a_s are UV-absorbance spectroscopy (UV-metric titration) [28–30], potentiometry (pH-metric titration) [30, 31], capillary electrophoresis [32, 33], and NMR spectroscopy [21], with NMR being the most time-consuming approach. Other, less popular p*K*_a_ measurement techniques include conductometry, HPLC, solubility or partition based estimations, calorimetry, fluorometry, and polarimetry [34]. UV-metric and pH-metric methods(Figure 3) of Sirius T3 are limited to measuring aqueous p*K*_a_ values between 2 and 12 due to limitations of the pH electrode used in these measurements. The pH-metric method relies on determining the stoichiometry of bound protons with respect to pH, calculated from volumetric titration with acid or base solutions. Accurate pH-metric measurements require high concentrations of analyte as well as analytically prepared acid/base stocks and analyte solutions. By contrast, UV-metric p*K*_a_ measurements rely on the differences in UV absorbance spectra of different protonation states, generally permitting lower concentrations of analyte to be used. The pH and UV absorbance of the analyte solution are monitored during titration.

**Figure 3.**
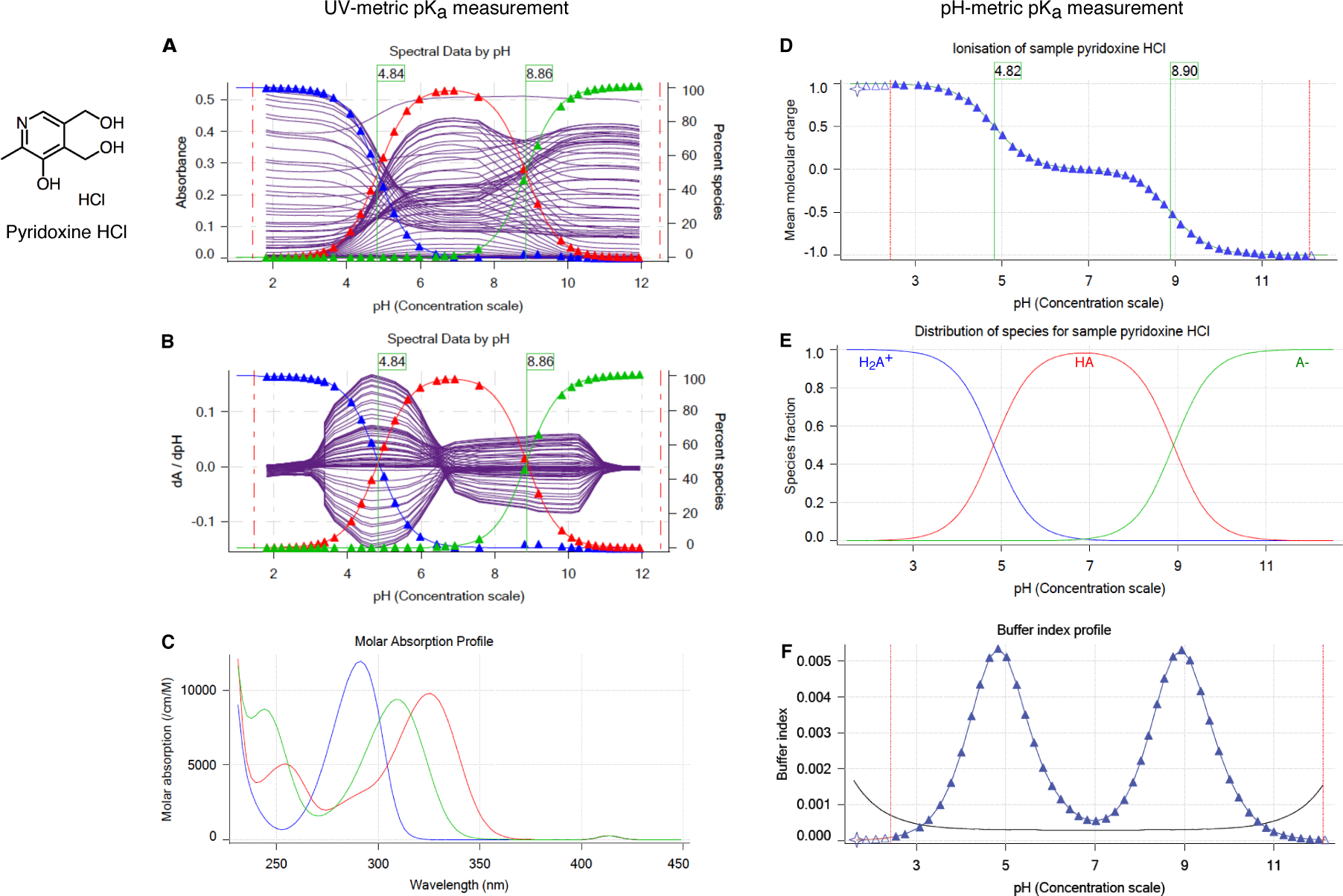
UV-metric (spectrophotometric) and pH-metric (potentiometric) p*K*_a_ measurements of pyridoxine HCl with Sirius T3. Spectrophotometic p*K*_a_ measurement (panels **A, B, C**) relies on differences in the UV absorbance spectra between microscopic protonation states to deconvolute the population of macrostate species as a function of pH. While highly sensitive (and therefore requiring a very low analyte concentration of ~ 50 μM), this approach can only resolve changes in protonation states for titratable sites near chromophores and cannot separate the populations of microstates that change in the same manner as a function of pH. (**A**) Multiwavelength UV absorbance vs pH. Purple lines represents absorbance at distinct wavelengths in UV region. (**B**) Derivative of multiwavelength absorbance with respect to pH (dA/dpH) vs pH is plotted with purple lines. In **A** and **B**, blue, red, and green triangles represent population of protonation states (from most protonated to least protonated) as calculated from a global fit to experimental UV absorbances for all pH values, while thin lines denote model fits that utilize the fitted model p*K*_a_s to compute populations. p*K*_a_ values (green fags) correspond to infection point of multiwavelength absorbance data where change in absorbance with respect to pH is maximum. (**C**) Molar absorption coefficients vs wavelength for each protonation state as resolved by TFA. **D, E, F** illustrate potentiometric p*K*_a_ measurement where molar addition of acid or base is tracked as pH is titrated. (**D**) Mean molecular charge vs pH. Mean molecular charge is calculated based on the model provided for the analyte: predicted number and nature of titratable sites (acid or base type), and number of counter ions present. p*K*_a_ values are calculated as infection points of charge vs pH plot. (**E**) Predicted macroscopic protonation state populations vs pH calculated based on p*K*_a_ values (H_2_A^+^: blue, HA: red, and A^-^: green) (**F**) Buffering index vs pH profile of water (grey solid line, theoretical) and the sample solution (blue triangles represent experimental data points). A higher concentration of analyte (~5 mM) is necessary for the potentiometric method than the spectrophotometric method in order to provide large enough buffering capacity signal above water for an accurate measurement.

Both UV-metric and pH-metric p*K*_a_ determination methods measure macroscopic p*K*_a_s for polyprotic molecules, which cannot be easily assigned to individual titration sites and underlying microstate populations in the absence of other experimental evidence that provides structural information, such as NMR (Figure 2). Macroscopic populations observed in these two methods are composed of different combinations of microstates depending on the principles of measurement technique. In potentiometric titrations, microstates with same total charge will be observed as one macrostate, while in spectrophotometric titrations, protonation sites remote from chromophores might be spectroscopically invisible, and macrostates will be formed from collections of microstates that manifest similar UV-absorbance spectra.

For UV-metric method to resolve populations of microstates, sufficiently different UV spectra between microstates and sufficiently non-overlapping change of populations with respect to pH are needed. However, relative tautomer populations of microstates with the same total charge do not depend on pH and stay constant while pH is titrated (Figure 2B), therefore they cannot be resolved by UV-metric method. The pH-metric method also cannot resolve microstates that have the same total charge as shown in Figure 2C.

Spectrophotometric p*K*_a_ determination is more sensitive than potentiometric determination, requiring low analyte concentrations (50–100 μM)—especially advantageous for compounds with low solubilities—but is only applicable to titration sites near chromophores. For protonation state changes to affect UV absorbance, a useful rule of thumb is that the protonation site should be a maximum of four heavy atoms away from the chromophore, which might consist of conjugated double bonds, carbonyl groups, aromatic rings, etc. Although potentiometric measurements do not suffer from the same observability limitations, higher analyte concentrations (~5 mM) are necessary for the analyte to provide sufficiently large enough buffering capacity signal above water to produce an accurate measurement. The accuracy of p*K*_a_s fit to potentiometric titrations can also be sensitive to errors in the estimated concentration of the analyte in the sample solution, while UV-metric titrations are insensitive to concentration errors. We therefore decided to adopt spectrophotometric measurements for collecting the experimental p*K*_a_ data for this challenge, and selected a compound set to ensure that all potential titration sites are in the vicinity of UV chromophores.

Here, we report on the selection of SAMPL6 p*K*_a_ challenge compounds, their macroscopic p*K*_a_ values measured by UV-metric titrations using a Sirius T3, as well as NMR-based microstate characterization of two SAMPL6 compounds with ambiguous protonation states associated with the observed macroscopic p*K*_a_s (SM07 and SM14). We discuss implications of the use of this experimental technique for the interpretation of p*K*_a_ data, and provide suggestions for future p*K*_a_ data collection efforts with the goal of evaluating or training computational p*K*_a_ predictions.

## Methods

### Compound selection and procurement

To select a set of small molecules focusing on the chemical space representative of kinase inhibitors for physicochemical property prediction challenges (p*K*_a_ and lipophilicity) we started from the kinase-targeted subclass of the ZINC15 chemical library [35] and applied a series of filtering and selection rules as depicted in Figure 4A. We focused on the availability "now" and reactivity "anodyne" subsets of ZINC15 in the first filtering step [http://zinc15.docking.org/subclasses/kinase/substances/subsets/now+anodyne/]. The "now" label indicates the compounds were availabile for immediate delivery, while the "anodyne" label excludes compounds matching filters that fag compounds with the potential for reactivity or pan-assay interference (PAINs) [36, 37].

**Figure 4.**
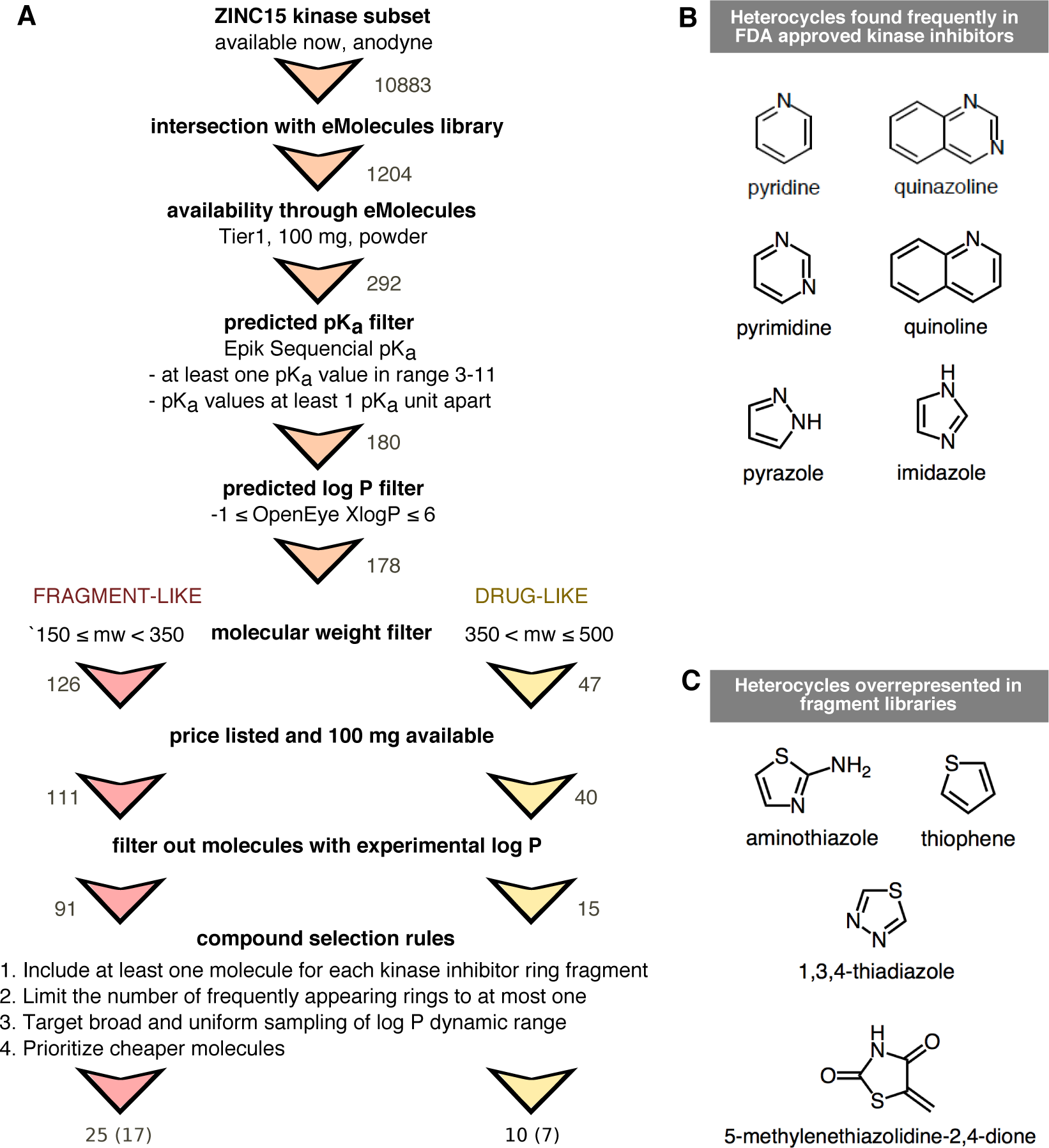
Compound selection for the SAMPL6 p*K*_a_ challenge, with the goal of running subsequent log *P*/log *D* challenges on the same compound set. (**A**) Flowchart of filtering steps for the selection of compounds that resemble kinase inhibitors and their fragments. Numbers next to arrows indicate the number of compounds remaining after each filtering step. A total of 25 fragment-like and 10 drug-like compounds were selected, out of which procurement and p*K*_a_ measurements for 17 fragment-like and 7 drug-like compounds were successful, respectively. (**B**) Frequent heterocycles found in FDA approved kinase inhibitors, as determined by Bemis-Murcko fragmentation into rings [49]. Black structures were represented in SAMPL6 set at least once. Compounds with piperazine and indazole (gray structures) could not be included in the challenge set due to library and selection limitations. (**C**) Structures of heterocycles that were overrepresented based on our compound selection workflow. We have limited the number of occurrences of these heterocycles to at most one.

Next, we identified resulting molecules that were also available for procurement through eMolecules [38] (free version, downloaded 1 June 2017), the supplier that would be used for procurement in this exercise. To find the intersection of ZINC15 kinase subset and eMolecules database, we matched molecules using their canonical isomeric SMILES strings, as computed via the OpenEye OEChem Toolkit (version 2017.Feb.1) [39].

To extract availability and price information from eMolecules, we queried using a list of SMILES (as reported in eMolecules database) of the intersection set. We further filtered the intersection set (1204 compounds) based on delivery time (Tier 1 suppliers, two-week delivery) and at least 100 mg availability in powder form (format: Supplier Standard Vial). We aimed to purchase 100 mg of each compound in powder form with at least 90% purity. We calculated 100 mg was enough for optimization and replicate experiments to measure p*K*_a_, log *P*, and solubility measurements with the Sirius T3. Each UV-metric and pH-metric p*K*_a_ measurement requires a minimum of 0.01 mg and 1.00 mg compound (solid or delivered via DMSO stock solution), respectively. log *P* and pH-dependent solubility measurements require around 2 mg and 10 mg of solid chemical, respectively.

### Filtering for predicted measurable p*K*_a_s and lack of experimental data

The Sirius T3 (Pion) instrument used to collect p*K*_a_ and log P/log *D* measurements requires a titratable group in the p*K*_a_ range of 2–12, so we aimed to select compounds with predicted p*K*_a_s in the range of 3-11 to allow a ~1 pK unit margin of error in p*K*_a_ predictions. p*K*_a_ predictions for compound selection were calculated using Epik (Schödinger) sequential p*K*_a_ prediction (scan) [40, 41] with target pH 7.0 and tautomerization allowed for generated states. We filtered out all compounds that did not have any predicted p*K*_a_s between 3–11, as well as compounds with two p*K*_a_ values predicted to be less than 1 p*K*_a_ unit apart in the hopes that individual p*K*_a_s of multiprotic compounds could be resolved with spectrophotometric p*K*_a_ measurements. With the goal of selecting compounds suitable for subsequent log *P* measurements, we eliminated compounds with OpenEye XlogP [42] values less than ‐1 or greater than 6. Subsets of compounds with molecular weights between 150-350 g/mol and 350–500 g/mol were selected for fragment-like and drug-like categories respectively. Compounds without available price or stock quantity information were eliminated. As the goal was to provide a blind challenge, compounds with publicly available experimental log *P* measurements were also removed. The sources we checked for publicly available experimental log *P* values were the following: DrugBank [43] (queried with eMolecules SMILES), ChemSpider [44] (queried by canonical isomeric SMILES), NCI Open Database August 2006 release [45], Enhanced NCI Database Browser [46] (queried with canonical isomeric SMILES), and PubChem [47] (queried with InChIKeys generated from canonical isomeric SMILES with NCI CACTUS Chemical Identifier Resolver [48]).

### Filtering for kinase inhibitor-like scaffolds

In order to include common scaffolds found in kinase inhibitors, we analyzed the frequency of rings found in FDA-approved kinase inhibitors via Bemis-Murcko fragmentation using OEMedChem Toolkit of OpenEye [49, 50]. Heterocycles found more than once in FDA-approved kinase inhibitors are shown in Figure 4B. In selecting 25 compounds for the fragment-like set and 10 compounds for the drug-like set, we prioritized including at least one example of each heterocycle, although we failed to find compounds with piperazine and indazole that satisfied all other selection criteria. We observed that certain heterocycles (shown in Figure 4C) were overrepresented based on our selection criteria; therefore, we limited the number of these structures in the SAMPL6 challenge set to at most one in each set. To achieve broad and uniform sampling of the measurable log *P* dynamic range, we segregated the molecules into bins of predicted XlogP values and selected compounds from each bin, prioritizing less expensive compounds.

### Filtering for UV chromophores

The presence of UV chromophores (absorbing in the 200–400 nm range) in close proximity to protonation sites is necessary for spectrophotometric p*K*_a_ measurements. To filter for molecules with UV chromophores, we looked at the substructure matches to the SMARTS pattern [`n,o,c`][`c,n,o`]`cc` which was considered the smallest unit of pi-conjugation that can constitute a UV chromophore. This SMARTS pattern describes extended conjugation systems comprised of four heavy atoms and composed of aromatic carbon, nitrogen, or oxygen, such as 1.3-butadiene, which possesses an absorption peak at 217 nm. Additionally, the final set of selected molecules was manually inspected to makes sure all potentially titratable groups were no more than four heavy atoms away from a UV chromophore.

25 fragment-like and 10 drug-like compounds were selected, out of which procurement of 28 was completed in time. p*K*_a_ measurements for 17 (SM01–SM17) and 7 (SM18–SM24) were successful, respectively. The resulting set of 24 small molecules constitute the SAMPL6 p*K*_a_ challenge set. For the other four compounds, UV-metric p*K*_a_ measurements show no detectable p*K*_a_s in the range of 2–12, so we decided not to include them in the SAMPL6 p*K*_a_ challenge. Experiments for these four compounds are not reported in this publication.

Python scripts used in the compound selection process are available from GitHub [https://github.com/choderalab/sampl6-physicochemical-properties]. Procurement details for each compound can be found in Supplementary Table 1. Chemical properties used in the selection of compounds are summarized in Supplementary Table 2.

### UV-metric p*K*_a_ measurements

Experimental p*K*_a_ measurements were collected using the spectrophotometric p*K*_a_ measurement method with a Sirius T3 automated titrator instrument (Pion) at 25°C and constant ionic strength. The Sirius T3 is equipped with an Ag/AgCl double-junction reference electrode to monitor pH, a dip probe attached to UV spectrophotometer, a stirrer, and automated volumetric titration capability. The Sirius T3 UV-metric p*K*_a_ measurement protocol measures the change in multi-wavelength absorbance in the UV region of the absorbance spectrum while the pH is titrated over pH 1.8–12.2 [28, 29]. UV absorbance data is collected from 160–760 nm while the 250–450 nm region is typically used for p*K*_a_ determinations. Subsequent global data analysis identifies pH-dependent populations of macrostates and fits one or more p*K*_a_ values to match this population with a pH-dependent model.

DMSO stock solutions of each compound with 10 mg/ml concentration were prepared by weighing 1 mg of powder chemical with a Sartorius Analytical Balance (Model: ME235P) and dissolving it in 100 μL DMSO (Dimethyl sulfoxide, Fisher Bioreagents, CAT: BP231-100, LOT: 116070, purity ≥ 99.7%). DMSO stock solutions were capped immediately to limit water absorption from the atmosphere due to the high hygroscopicity of DMSO and sonicated for 5–10 minutes in a water bath sonicator at room temperature to ensure proper dissolution. These DMSO stock solutions were stored at room temperature up to two weeks in capped glass vials. 10 mg/ml DMSO solutions were used as stock solutions for the preparation of three replicate samples for the independent titrations. For each experiment, 1–5 μL of 10 mg/ml DMSO stock solution was delivered to a 4 mL Sirius T3 glass sample vial with an electronic micropipette (Rainin EDP3 LTS 1–10 μL). The volume of delivered DMSO stock solution, which determines the sample concentration following dilution by the Sirius T3, is optimized individually for each compound to achieve sufficient but not saturated absorbance signal (targeting 0.5–1.0 AU) in the linear response region. Another limiting factor for sample concentration was ensuring that the compound remains soluble throughout the entire pH titration range. An aliquot of 25 μL of mid-range buffer (14.7 mM K_2_HPO_4_ and 0.15 M KCl in H_2_O) was added to each sample, transferred with a micropipette (Rainin EDP3 LTS 10–100 μL) to provide enough buffering capacity in middle pH ranges so that pH could be controlled incrementally throughout the titration.

pH is temperature and ionic-strength dependent. A peltier device on the Sirius T3 kept the analyte solution at 25.0 ± 0.5 °C throughout the titration. Sample ionic strength was adjusted by dilution in 1.5 mL ionic strength-adjusted water (ISA water ≡ 0.15 M KCl in H_2_O) by the Sirius T3. Analyte dilution, mixing, acid/base titration, and measurement of UV absorbance was automated by the Sirius T3 UV-metric p*K*_a_ measurement protocol. The pH was titrated between pH 1.8 and 12.2 via the addition of acid (0.5 M HCl) and base (0.5 M KOH), targeting 0.2 pH steps between UV absorbance spectrum measurements. Titrations were performed under argon flow on the surface of the sample solution to limit the absorption of carbon dioxide from air, which can alter the sample pH to a measurable degree. To fully capture all sources of experimental variability, instead of performing three sequential pH titrations on the same sample solution, three replicate samples (prepared from the same DMSO stock solution) were subjected to one round of pH titration each. Although this choice reduced throughput and increased analyte consumption, it limited the dilution of the analyte during multiple titrations, resulting in stronger absorbance signal for p*K*_a_ fitting. Under circumstances where analyte is scarce, it is also possible to do three sequential titrations using the same sample to limit consumption when the loss of accuracy is acceptable.

Visual inspection of the sample solutions after titration and inspection of the pH-dependent absorbance shift in the 500–600 nm region of the UV spectra was used to verify no detectable precipitation occurred during the course of the measurement. Increased absorbance in the 500–600 nm region of the UV spectra is associated with scattering of longer wavelengths of light in the presence of colloidal aggregates. For each analyte, we optimized analyte concentration, direction of the titration, and pH titration range in order to maintain solubility over the entire experiment. The titration direction was specified so that each titration would start from the pH where the compound is most soluble: low-to-high pH for bases and high-to-low pH for acids. While UV-metric p*K*_a_ measurements can be performed with analyte concentrations as low as 50 μM (although this depends on the absorbance properties of the analyte), some compounds may yet not be soluble at these low concentrations throughout the pH range of the titration. As the sample is titrated through a wide range of pH values, it is likely that low-solubility ionization states—such as neutral and zwitterionic states—will also be populated, limiting the highest analyte concentration that can be titrated without encountering solubility issues. For compounds with insufficient solubility to accurately determine a p*K*_a_ value directly in a UV-metric titration, a cosolvent protocol was used, as described in the next section (**UV-metric p*K*_a_ measurement with cosolvent**).

Two Sirius T3 computer programs—Sirius T3 Control v1.1.3.0 and Sirius T3 Refine v1.1.3.0—were used to execute measurement protocols and analyze pH-dependent multiwavelength spectra, respectively. Protonation state changes at titratable sites near chromophores will modulate the UV-absorbance spectra of these chromophores, allowing populations of distinct UV-active species to be resolved as a function of pH. To do this, basis spectra are identified and populations extracted via TFA analysis of the pH-dependent multi-wavelength absorbance [29]. When fitting the absorbance data to a titratable molecule model to estimate p*K*_a_s, we selected the minimum number of p*K*_a_s sufficient to provide a high-quality fit between experimental and modeled data based on visual inspection of pH-dependent populations.

This method is capable of measuring p*K*_a_ values between 2–12 when titratable groups are at most 4–5 heavy atoms away from chromophores such that a change in protonation state alters the absorbance spectrum of the chromophore. We selected compounds where titratable groups are close to potential chromophores (generally aromatic ring systems), but the possibility exists that our experiments did not detect protonation state changes of titratable groups distal from UV chromophores.

### Cosolvent UV-metric p*K*_a_ measurements of molecules with poor aqueous solubilities

If analytes are not sufficiently soluble during the titration, p*K*_a_ values cannot be accurately determined via aqueous titration directly. If precipitation occurs, the UV-absorbance signal from pH-dependent precipitate formation cannot be differentiated from the pH-dependent signal of soluble microstate species. For compounds with low aqueous solubility, p*K*_a_ values were estimated from multiple apparent p*K*_a_ measurements performed in ISA methanol:ISA water cosolvent solutions with various mole fractions, from which the p*K*_a_ at 0% methanol (100% ISA water) can be extrapolated. This method is referred to as a UV-metric p_s_K_a_ measurement in the Sirius T3 Manual [51]. p_s_K_a_ value is the apparent p*K*_a_ value measured in the presence of a cosolvent.

The cosolvent spectrophotometric p*K*_a_ measurement protocol was very similar to the standard aqueous UV-metric p*K*_a_ measurement protocol, with the following differences: titrations were performed in typically in 30%, 40%, and 50% mixtures of ISA methanol:ISA water by volume to measure apparent p*K*_a_ values (p_s_K_a_) in these mixtures. Yasuda-Shedlovsky extrapolation [52, 53] was subsequently used to estimate the p*K*_a_ value at 0% cosolvent (Figure 5) [31, 54, 55].

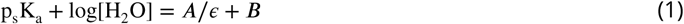

**Figure 5.**
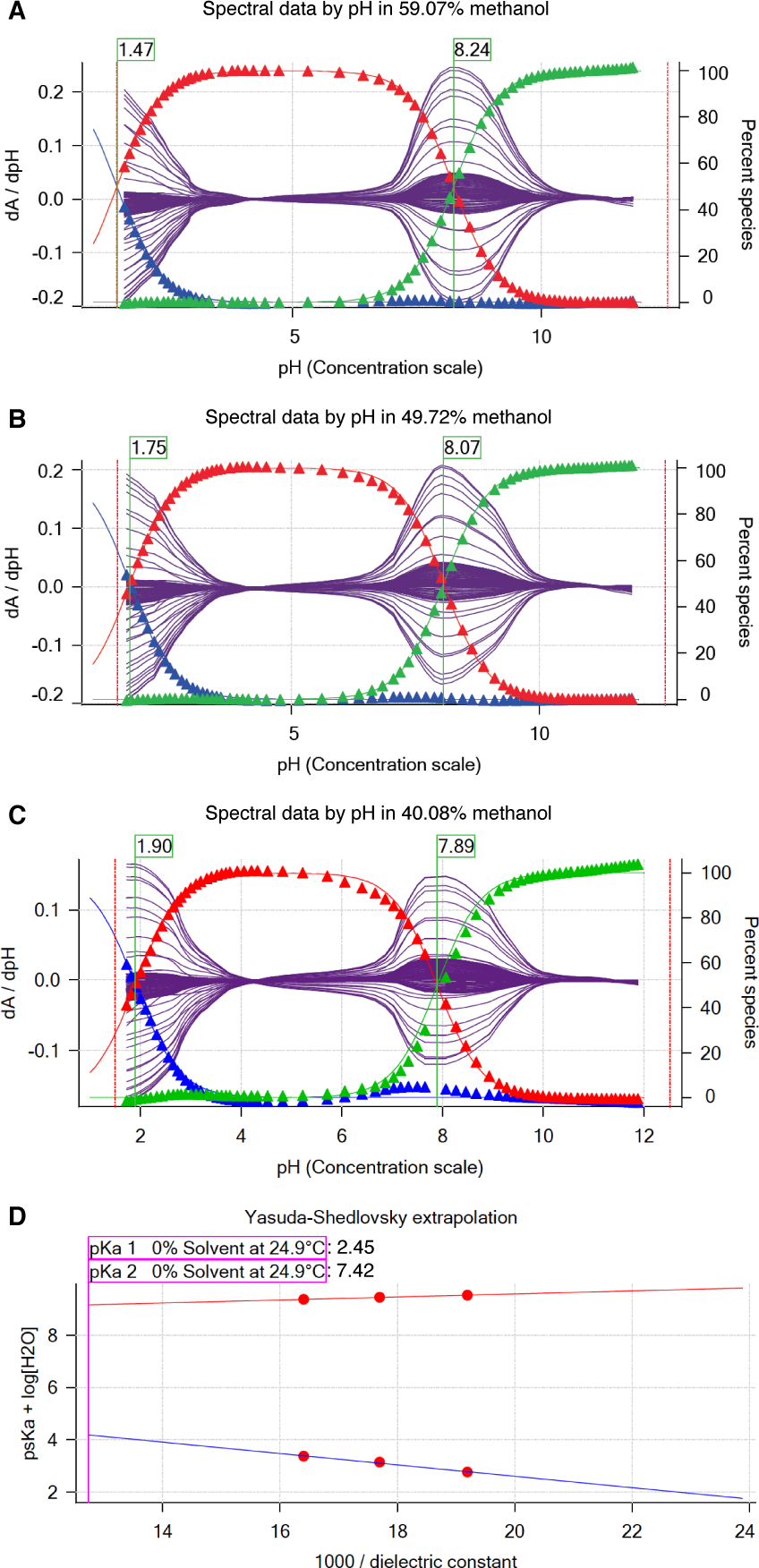
Determination of SM22 p*K*_a_ values with cosolvent method and Yasuda-Shedlovsky extrapolation. **A, B**, and **C** show p_s_*K*_a_ of SM22 determined at various methanol concentrations: 59.07%, 49.72%, 40.08% by weight. Purple solid lines indicate the derivative of the absorbance signal with respect to pH i/s pH at multiple wavelengths. p_s_*K*_a_ values (green fags) were determined by Sirius T3 Refine Software. Blue, red, and green triangles show relative populations of macroscopic protonation states with respect to pH calculated from the experimental data. Notice that as cosolvent concentration increases, p_s_*K*_a1_ decreases from 1.90 to 1.47 and p_s_*K*_a2_ increases from 7.84 to 8.24. **D** Yasuda-Shedlovsky extrapolation plot for SM22. Red datapoints correspond to p_s_*K*_a_ determined at various cosolvent ratios. Based on linear fitting to *p*_*s*_***K***_*a*_ + *log*\H_2_O] vs 1/*ϵ*, p*K*_a1_ and p*K*_a2_ in 0% cosolvent (aqueous solution) was determined as 2.45 and 7.42, respectively. R^2^ values of linear fits are both 0.99. The slope of Yasuda-Shedlovsky extrapolation shows if the observed titration has acidic (positive slope) or basic (negative slope) character dominantly, although this is an macroscopic observation and should not be relied on for annotation of p*K*_a_s to functional groups (microscopic p*K*_a_s).

Yasuda-Shedlovsky extrapolation relies on the linear correlation between p_s_K_a_ + log[H_2_O] and the reciprocal dielectric constant of the cosolvent mixture (1/*ϵ*). In Eq. 1, A and B are the slope and intercept of the line fitted to experimental datapoints. Depending on the solubility requirements of the analyte, the methanol ratio of the cosolvent mixtures was adjusted. We designed the experiments to have at least 5% cosolvent ratio difference between datapoints and no more than 60% methanol content. Calculation of the Yasuda-Shedlovsky extrapolation was performed by the Sirius T3 software using at least 3 p_s_*K*_a_ values measured in different ratios of methanol:water. Addition of methanol (80%, 0.15 M KCl) was controlled by the instrument before each titration. Three consecutive pH titrations at different methanol concentrations were performed using the same sample solution. In addition, three replicate measurements with independent samples (prepared from the same DMSO stock) were collected.

### Calculation of uncertainty in p*K*_a_ measurements

Experimental uncertainties were reported as the standard error of the mean (SEM) of three replicate p*K*_a_ measurements. The standard error of the mean (SEM) was estimated as

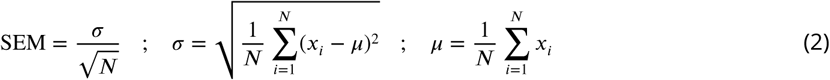

where *σ* denotes the sample standard deviation and *μ* denotes the sample mean. *x*_*i*_ are observations and N is the number of observations.

Since the Sirius T3 software reports p*K*_a_ values to only two decimal places, we have reported the SEM as 0.01 in cases where SEM values calculated from 3 replicates were lower than 0.01. SEM calculated from replicate measurements were found to be larger than non-linear fit error reported by the Sirius T3 Refine Software from UV-absorbance model fit of a single experiment, thus leading us to believe that running replicate measurements and reporting mean and SEM of p*K*_a_ measurements is better for capturing all sources of experimental uncertainty. Notably, for UV-metric measurements, the measured p*K*_a_ values should be insensitive to final analyte concentration and any uncertainty in the exact analyte concentration of the original DMSO stock solution, justifying the use of the same stock solution (rather than independently prepared stock solutions) for multiple replicates.

### Quality control for chemicals

Compound purity was assessed by LC-MS using an Agilent HPLC 1200 Series equipped with auto-sampler, UV diode array detector, and a Quadrupole MS detector 6140. ChemStation version C01.07SR2 was used to analyze LC & LC/MS. An Ascentis Express C18 column (3.0 x 100 mm, 2.7 μm) was used, with column temperature set at 45° C.

- Mobile phase A: 2 mM ammonium formate (pH = 3.5) aqueous
- Mobile phase B: 2 mM ammonium formate in 90:10 acetonitrile:water (pH = 3.5)
- Flow rate : 0.75 ml/min
- Gradient: Starting with 10% B to 95% B in 10 minutes then hold at 95% B for 5 minutes.
- Post run length: 5 minutes
- Mass condition: ESI positive and negative mode
- Capillary voltage: 3000 V
- Drying gas flow: 12 ml/min
- Nebulizer pressure: 35 psi
- Drying temperature: 350°C
- Mass range: 5-1350 Da; Fragmentor: 70; Threshold: 100

The percent area for the primary peak is calculated based on the area of the peak divided by the total area of all peaks. The percent area of the primary peak is reported as an estimate of sample purity. The purity of primary LC peak was checked by ChemStation software with threshold 995, to check that there is no significant impurity underneath the main peak.

### NMR determination of protonation microstates

In general, the chemical shifts of nuclear species observed in nuclear magnetic resonance (NMR) spectra report on and are very sensitive to the chemical environment. Consequently, small changes in chemical environment, such as the protonation events described in this work, are manifest as changes in the chemical shift(s) of the nuclei. If perturbation occurs at a rate which is fast on the NMR timescale (*fast exchange*), an average chemical shift is observed. This phenomena has been exploited and utilized as a probe for determining the order of protonation for molecules with more than one titratable site [56]. In some cases, direct observation of the titrated nuclei can be difficult, for example nitrogen and oxygen, due to sample limitations and/or low natural abundance of the NMR active nuclei (0.37% for ^15^N and 0.038% for ^17^O)—amongst other factors. In these situations, chemical shifts changes of the so-called “reporter” NMR nuclei—^1^H, ^31^P, or ^13^C nuclei, which are directly attached to or are a few bonds away from the titrated nuclei—have been utilized as the probe for NMR-pH titrations [21, 57, 58]. This approach is advantageous since the sensitive NMR nuclides (^1^H and ^31^P) are observed. In addition, ^31^P and ^13^C offer large spectral widths of ~300 ppm and ~200 ppm, respectively, which minimize peak overlap.

However, reporter nuclei chemical shifts provide indirect information subject to interpretation. In complex systems with multiple titratable groups, such analysis will be complicated due to a cumulative effect of these groups on the reporter nuclide due to their close proximity or the resonance observed in aromatic systems. In contrast, direct observation of the titratable nuclide where possible, affords a more straight-forward approach to studying the protonation events. In this study, the chemical shifts of the titratable nitrogen nuclei were observed using the ^1^H-^15^N-HMBC (Heteronuclear Multiple-Bond Correlation) experiments — a method that affords the observation of ^15^N chemical shifts while leveraging the sensitivity accrued from the high abundance the ^1^H nuclide.

The structures of samples SM07 and SM14 were assigned via a suite of NMR experiments, which included ^1^H NMR, ^13^C NMR, homonuclear correlated spectroscopy (^1^H-^1^H COSY), heteronuclear single quantum coherence (^1^H-^13^C HSQC), ^13^C heteronuclear multiple-bond correlation (^1^H-^13^C-HMBC) and ^15^N heteronuclear multiple-bond correlation (^1^H-^15^N-HMBC)—see SI. All NMR data used in this analysis were acquired on a Bruker 500 MHz spectrometer equipped with a 5 mm TCI CryoProbe™ Prodigy at 298 K. The poor solubility of the analytes precluded analysis in water and thus water-*d*_2_/methanol-*d*_4_ mixture and acetonitrile-*d*_3_ were used as solvents. The basic sites were then determined by titration of the appropriate solutions of the samples with equivalent amounts of deutero-trif uoroacetic acid (TFA-d) solution.

### SM07

5.8 mg of SM07 was dissolved in 600 μL of methanol-*d*_4_:water-*d*_2_ (2:1 v/v ratio). A 9% v/v TFA-d solution in water-d2 was prepared, such that each 20 μL volume contained approximately 1 equivalent of TFA-*d* with respect to the base. The SM07 solution was then titrated with the TFA-*d* solution at 0.5, 1.0, 1.5, and 5.0 equivalents with ^1^H-^15^N HMBC spectra (optimized for 5 Hz) acquired after each TFA addition. A reference ^1^H-^15^N HMBC experiment was first acquired on the SM07 solution prior to commencement of the titration.

### SM14

5.5 mg of SM14 was dissolved in 600 μL of acetonitrile-*d*_3_. A 10% v/v TFA-d solution in acetonitrile-*d*_3_ was prepared, 20 μL of which corresponds to 1 equivalent of TFA-d with respect to the base. Further 1:10 dilution of the TFA-d solution in acetonitrile-*d*_3_, allowed measurement of 0.1 equivalent of TFA-d per 20 μL of solution. The SM14 solution was then titrated with the TFA-d solutions at 0.0, 0.5, 1.0, 1.1, 1.2, 1.3, 1.5, 1.8, 2.0, 2.1, 2.6, 5.1, and 10.1 equivalents. The chemical shift changes were monitored by the acquisition of ^1^H-^15^N HMBC spectra (optimized for 5 Hz) after each TFA addition.

## Results

### Spectrophotometric p*K*_a_ measurements

Spectrophotometrically-determined p*K*_a_ values for all molecules from the SAMPL6 p*K*_a_ challenge are shown in Figure 6 and Table 1. The protocol used—cosolvent or aqueous UV-metric titration—is indicated in Table 1 together with SEM of each reported measurement. Out of 24 molecules successfully assayed, five molecules have two resolvable p*K*_a_ values, while one has three resolvable p*K*_a_ values within the measurable p*K*_a_ range of 2-12. The SEM of reported p*K*_a_ measurements is low, with the largest uncertainty reported being 0.04 pK units (p*K*_a1_ of SM06 and pK_a3_ of SM18). Individual replicate measurements can be found in Supplementary Table 3. Reports generated for each p*K*_a_ measurement by the Sirius T3 Refine software can also be found in the Supplementary Information. Experimental p*K*_a_ values for nearly all compounds with multiple resolvable p*K*_a_s are well-separated (more than 4 p*K*_a_ units), except for SM14 and SM18.

**Figure 6.**
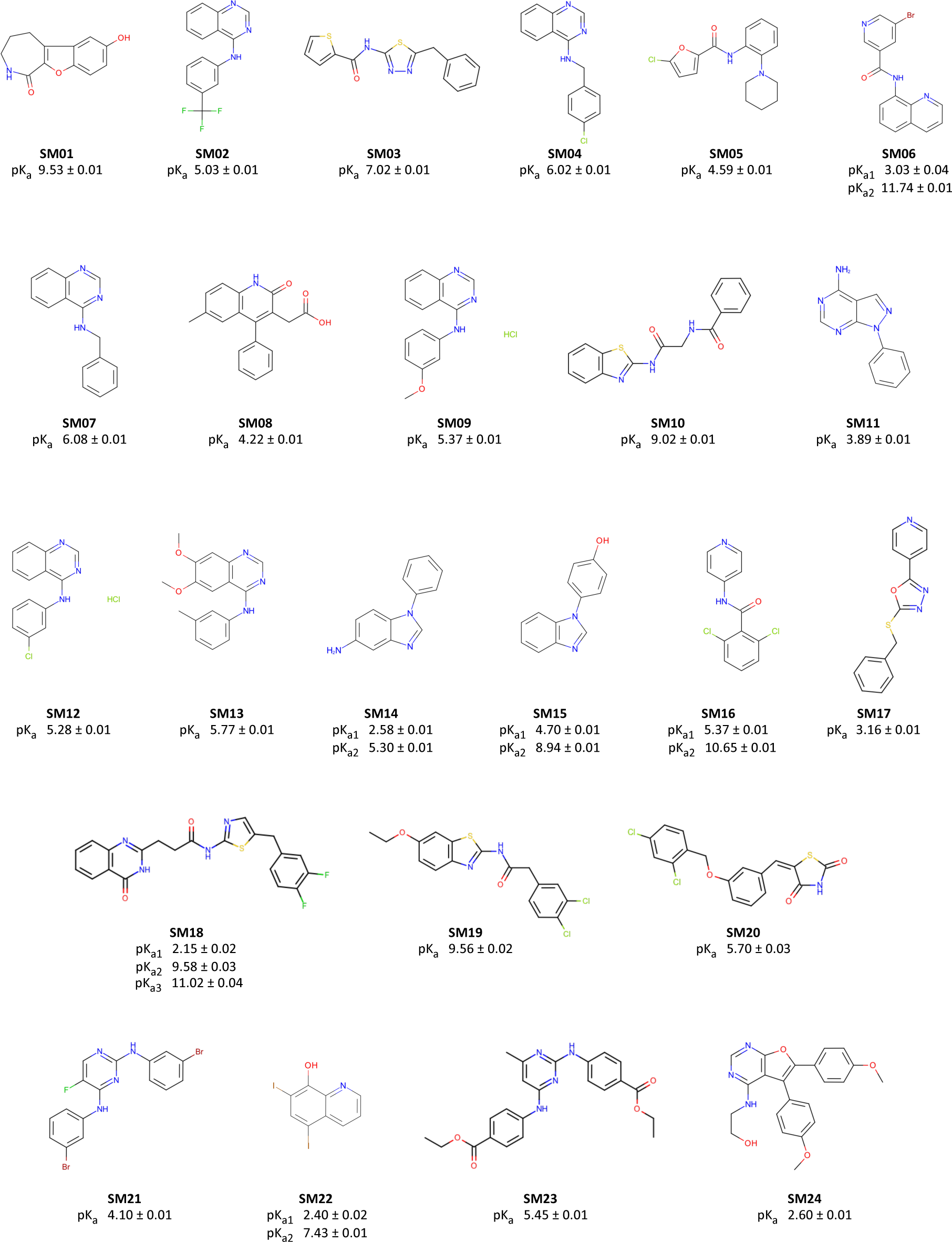
Molecules used in the SAMPL6 p*K*_a_ challenge. Experimental UV-metric p*K*_a_ measurements were performed for these 24 molecules and discernable macroscopic p*K*_a_s are reported. Uncertainties are expressed as the standard error of the mean (SEM) of three independent measurements. We depicted neutral states of the molecules as sites of protonation were not determined by UV-metric methods. 2D structures were created with OpenEye OEDepict Toolkit [59]. Canonical isomeric SMILES of molecules in this figure and p*K*_a_ values measured in replicate experiments can be found in Table SI 1 and Table SI 3, respectively.

**Table 1.** Experimental p*K*_a_s of SAMPL6 compounds. Spectrophotometric p*K*_a_ measurements were performed with two assay types based on aqueous solubility of analytes. "UV-metric p*K*_a_" assay indicates spectrophotometric p*K*_a_ measurements done with Sirius T3 in ISA water. "UV-metric p*K*_a_ with cosolvent" assay refers to p*K*_a_ determination by Yasuda-Shedlovsky extrapolation from p_s_*K*_a_ measurements in various ratios of ISA methanol:water mixtures. Triplicate measurements were performed at 25.0 ± 0.5° C and in the presence of approximately 150 mM KCl to adjust ionic strength.

| Molecule ID | $pK_{a1}$ | $pK_{a2}$ | $pK_{a3}$ | Assay Type |
| --- | --- | --- | --- | --- |
| SM01 | $9.53 \pm 0.01$ | | | UV-metric $pK_a$ |
| SM02 | $5.03 \pm 0.01$ | | | UV-metric $pK_a$ with cosolvent |
| SM03 | $7.02 \pm 0.01$ | | | UV-metric $pK_a$ with cosolvent |
| SM04 | $6.02 \pm 0.01$ | | | UV-metric $pK_a$ |
| SM05 | $4.59 \pm 0.01$ | | | UV-metric $pK_a$ with cosolvent |
| SM06 | $3.03 \pm 0.04$ | $11.74 \pm 0.01$ | | UV-metric $pK_a$ |
| SM07 | $6.08 \pm 0.01$ | | | UV-metric $pK_a$ |
| SM08 | $4.22 \pm 0.01$ | | | UV-metric $pK_a$ |
| SM09 | $5.37 \pm 0.01$ | | | UV-metric $pK_a$ with cosolvent |
| SM10 | $9.02 \pm 0.01$ | | | UV-metric $pK_a$ with cosolvent |
| SM11 | $3.89 \pm 0.01$ | | | UV-metric $pK_a$ |
| SM12 | $5.28 \pm 0.01$ | | | UV-metric $pK_a$ |
| SM13 | $5.77 \pm 0.01$ | | | UV-metric $pK_a$ |
| SM14 | $2.58 \pm 0.01$ | $5.30 \pm 0.01$ | | UV-metric $pK_a$ |
| SM15 | $4.70 \pm 0.01$ | $8.94 \pm 0.01$ | | UV-metric $pK_a$ |
| SM16 | $5.37 \pm 0.01$ | $10.65 \pm 0.01$ | | UV-metric $pK_a$ |
| SM17 | $3.16 \pm 0.01$ | | | UV-metric $pK_a$ |
| SM18 | $2.15 \pm 0.02$ | $9.58 \pm 0.03$ | $11.02 \pm 0.04$ | UV-metric $pK_a$ with cosolvent |
| SM19 | $9.56 \pm 0.02$ | | | UV-metric $pK_a$ with cosolvent |
| SM20 | $5.70 \pm 0.03$ | | | UV-metric $pK_a$ with cosolvent |
| SM21 | $4.10 \pm 0.01$ | | | UV-metric $pK_a$ with cosolvent |
| SM22 | $2.40 \pm 0.02$ | $7.43 \pm 0.01$ | | UV-metric $pK_a$ with cosolvent |
| SM23 | $5.45 \pm 0.01$ | | | UV-metric $pK_a$ with cosolvent |
| SM24 | $2.60 \pm 0.01$ | | | UV-metric $pK_a$ with cosolvent |

<sup>1</sup> $pK_a$ values are reported as mean ± SEM of three replicates.

### Impact of cosolvent to UV-metric p*K*_a_ measurements

For molecules with insufficient aqueous solubilities throughout the titration range (pH 2-12), we resorted to cosolvent UV-metric p*K*_a_ measurements, with methanol used as cosolvent. To confirm that cosolvent UV-metric p*K*_a_ measurements led to indistinguishable results compared to aqueous UV-metric measurements, we collected p*K*_a_ values of 12 highly soluble SAMPL6 compounds—as well as pyridoxine—using both cosolvent and aqueous methods. Correlation analysis of p*K*_a_ values determined with both methods demonstrated that using methanol as cosolvent and determining aqueous p*K*_a_s via Yasuda-Shedlovsky extrapolation did not result in significant bias (Figure 7), since 95% CI for mean deviation (MD) between two measurements includes zero. Means and standard errors of UV-metric p*K*_a_ measurements with and without cosolvent are provided in Supplementary Table 5. p*K*_a_ measurement results of individual replicate measurements with and without cosolvent can be found in Supplementary Table 4.

**Figure 7.**
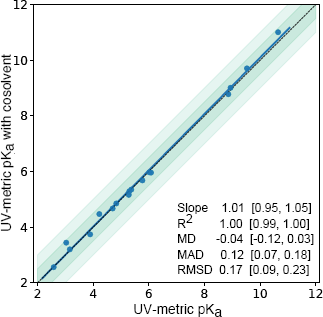
p*K*_a_ measurements with UV-metric method with cosolvent and UV-metric method in water show good correlation. 17 p*K*_a_ values (blue marks) of 13 chemicals were measured with both UV-metric p*K*_a_ method in water and UV-metric p*K*_a_ method with methanol as cosolvent (Yasuda-Shedlovsky extrapolation to 0% methanol). Dashed black line has slope of 1, representing perfect correlation. Dark and light green shaded areas indicate ±0.5 and ±1.0 p*K*_a_ unit difference regions, respectively. Error bars are plotted as the SEM of replicate measurements, although they are not visible since the largest SEM is 0.04. MD: Mean difference, MAD: Mean absolute deviation, RMSD: Root-mean-square deviation. Confidence intervals (reported in brackets) report the 95%ile CI calculated over 10 000 bootstrap samples. Experimental data used in this plot is reported in Supplementary Table 4.

### Purity of SAMPL6 compounds

LC-MS based purity measurements showed that powder stocks of 23 of the SAMPL6 p*K*_a_ challenge compounds were >90% pure, while purity of SM22 was 87%—the lowest in the set (Supplementary Table 6). Additionally, molecular weights detected by LC-MS method were consistent with those reported in eMolecules, as well as supplier-reported molecular weights, when provided. It is recommended by Sirius/Pion technical specialists to use compounds with ~90% purity to minimize the impact on high-accuracy p*K*_a_ measurements. Impurities with no UV-chromophore, or elute too late in LC may not be detected with this method, although chances are small. The peak purity check of primary peak can detect the presence of a large impurity underneath the main peak, but if the UV spectrum of the impurity is exactly same with analyte in the main peak, it may not be resolved. HPLC UV detector’s wavelength inaccuracy is <1%. Mass inaccuracy of MS instrument is ~0.13 um within the calibrated mass range in the scan mode.

### Characterization of SM07 microstates with NMR

^15^N Chemical shifts (ppm, referenced to external liquid ammonia at 0 ppm) for N-8, N-10 and N-12—measured from the ^1^H-^15^N HMBC experiments—were plotted against the titrated TFA-d equivalents (0.0, 0.5, 1.0, 1.5, and 5.0 equivalents) (Figure 8 A). A large upfield shift of ~82 ppm is observed for N-12. The initial linear relationship between chemical shift and TFA equivalents, shown in Figure 8A for N-12, is expected for strong monoprotic bases—as is the case for SM07. The large upfield chemical shift change (82 ppm) is consistent with a charge delocalization as shown in the resonance structures in Figure 8A. Further evidence for this delocalization is observed for N-8, which exhibited a downfield chemical shift change of ~28 ppm compared to just ~1.5 ppm for N-10. Titration of SM07 with more than 1 equivalent of TFA-*d* did not result in further significant chemical shift changes—establishing that SM07 is a monoprotic base.

**Figure 8.**
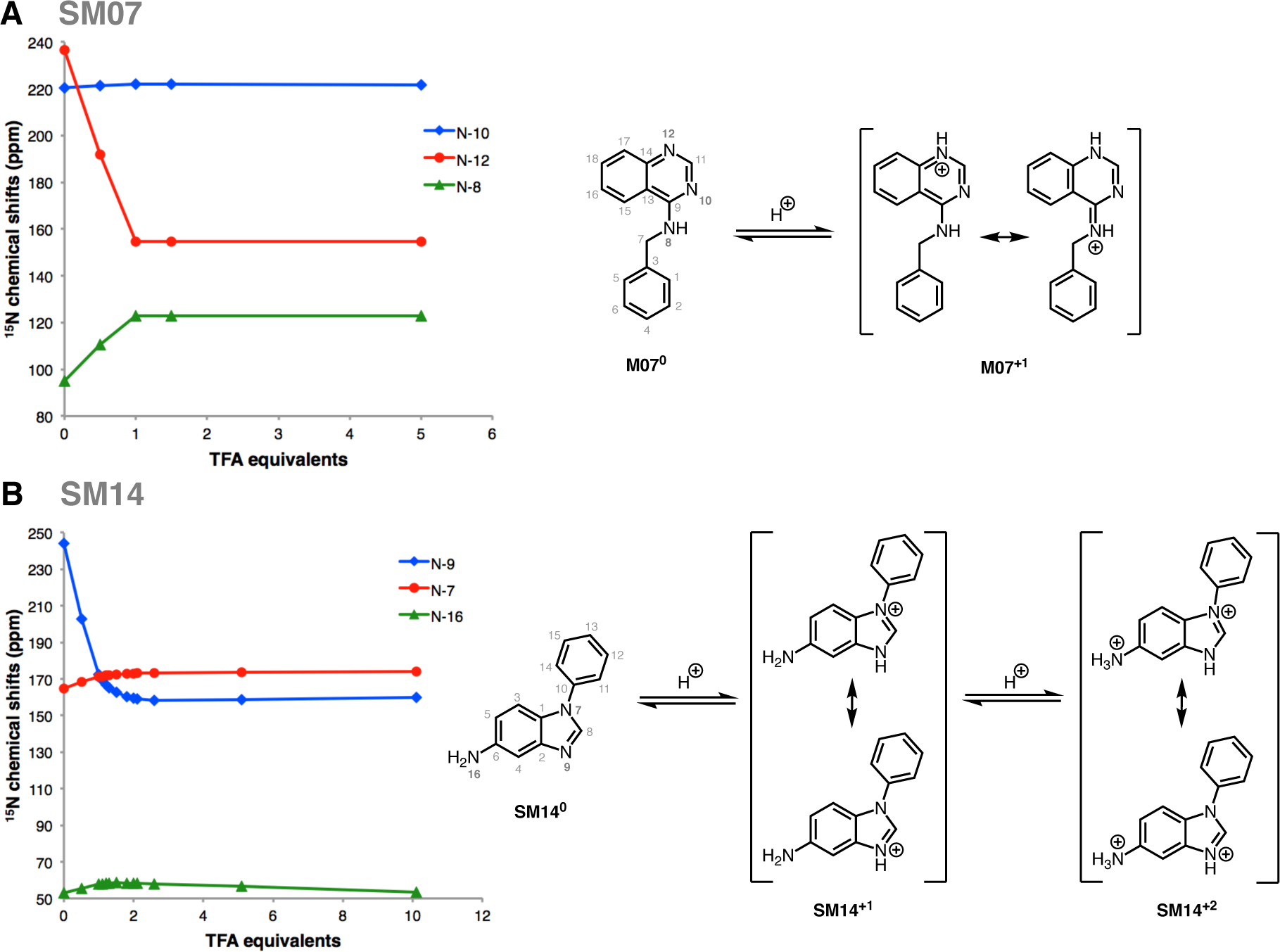
Dominant protonation microstates of SM07 and SM14 characterized by NMR. (**A**) Sequence of protonation sites of SM07 were determined by^1^ H-^15^N HMBC experiments in 1:2 water:methanol mixture. *Left:* The plot of ^15^N chemical shifts oftheN-10, N-12, and N-8 resonances of SM07 vs titrated TFA-*d* equivalents, showing the mono-protonation of N-12 as evidenced by its large upfield chemical shifts change. Acidity of the medium increased as more equivalents of TFA-*d* were added. Electronic effects due to protonation of N-12 caused downfield chemical shift change of N-10 and N-8 between 0-1 equivalents of TFA-*d*. *Right:* NMR-based model of the order of dominant protonation states for SM07. The protonation event was only observed at N-12. Microstates shown in the figure are the most likely contributors to the UV-metric pKa of 6.08 ± 0.01. (**B**) Sequence of protonation sites of SM14 were determined by^1^ H-^15^N HMBC experiments in acetonitrile. *Left:* The plot of^15^N chemical shifts of N-9, N-7, and N-16 of SM14 vs titrations of TFA-*d* equivalents, showing two sequential protonation events. The first protonation occured at N-9; a large upfield chemical shift change of 71.6 ppm was seen between 0-1 equivalents of TFA-*d*. Downfield chemical shift changes observed for N-7 and N-19 in this region were due the electronic effect from the protonation of N-9. N-16 also exhibited a small upfield chemical shift change of 4.4 ppm between 2.5-10 equivalents of TFA-d, which indicated N-16 as the second site of protonation. *Right:* NMR based model of the order of dominant protonation states for SM14, showing two sequential protonation events. Also, two pKa values were detected with UV-metric pKa measurements for SM14. Assuming that the sequence of protonation events will be conserved between water and acetonitrile solvents, SM14^0^ and SM14+^1^ microstates shown in the figure are the major contributors to the UV-metric pKa value 5.30 ± 0.01. SM14^+1^ and SM14^+2^ microstates shown in the figure are the major pair of microstates contributing to the UV-metric p*K*_a_ value 2.58 ± 0.01. There could be minor microstates with very low populations that could not be distinguished in these NMR experiments.

### Characterization of SM14 microstates with NMR

Determining the protonation sites for SM14, which has p*K*_a_ values of 2.58 and 5.30 (Table 1), was more challenging due to multiple possible resonance structures in the mono- and di-protonated states. We noticed that the water/methanol co-solvent exhibited strong solvent effects, which complicated the data interpretation for SM14. For instance, titration of SM14 in methanol/water (Figure SI 36) showed incomplete protonation of N-9 even after 5 equivalents of TFA-*d* were added. This observation is consistent with UV-metric p_s_*K*_a_ measurements done in the presence of methanol as cosolvent, where both p_s_*K*_a_ values were decreasing as the percentage of methanol was increased, making observation of these protonation states more difficult. Thus the utilization of an aprotic solvent was necessary for unambiguous interpretation of the data.

Due to the problem just delineated for the methanol/water cosolvent, acetonitrile-*d*_3_ was selected as our solvent of choice. Titration of SM14 (5.5 mg) with up to 10 equivalents of TFA-*d* in acetonitrile-*d*_3_ (0.0, 0.5, 1.0, 1.1, 1.2, 1.3, 1.5, 1.8, 2.0, 2.1, 2.6, 5.1, and 10.1 equivalents), provided a much clearer picture of its protonation states (Figure 8 B). N-9, with the large upfield chemical shift change ~72 ppm at 1 equivalent of TFA-*d*, clearly is the site of first protonation. Concurrently, the downfield chemical shift changes were observed for N-7 (Δ*δ* ≈ 6.5) and N-16 (Δ*δ* ≈ 5) that can be attributed to electronic effects rather than a direct protonation. The large upfield shift for N-9 indicates this to be the site of first protonation; complete protonation was attained at roughly 2.5 equivalents of TFA-*d*, suggesting that SM14 is a weak base under these experimental conditions. Following the protonation of N-9, a second protonation event occurs at N-16 nitrogen as evident by the upfield chemical shift change observed for N-16. However, a continuous change in the chemical shift of N-16 even after addition of 10 equivalents of TFA-*d* indicates that this protonation event is incomplete but provides evidence for N-16 being the second protonation site. This observation is consistent with N-16 being even a weaker base than N-9, which is expected of the aniline-type amines. Other notable observations were the slight downfield chemical shift changes for N-7 and N-9, during the second protonation event. These changes were attributed to electronic effects from the protonation of N-16.

## Discussion

### Effect of sample preparation and cosolvents in UV-metric measurements

Samples for UV-metric p*K*_a_ measurements were prepared by dilution of up to 5 μL DMSO stock solution of analyte in 1.5 mL ISA water, which results in the presence of ~0.3% DMSO during titration, which is presumed to have a negligible effect on p*K*_a_ measurements. For UV-metric or pH-metric measurements, it is possible to prepare samples without DMSO, but it is difficult to prepare samples by weighing extremely low amounts of solid stocks (in the order of 0.01–0.10 mg) to target 50 μM analyte concentrations, even with an analytical balance. For experimental throughput, we therefore preferred using DMSO stock solutions. Another advantage of starting from DMSO stock solutions is that it helps to overcome kinetic solubility problems of analytes.

A lower analyte concentration is needed for spectrophotometric p*K*_a_ measurement than potentiometric method. With spectrophotometric method, very dilute analyte solutions as low as 10^‒5^ – 10^‒6^ M can be used [28] with strength of the UV signal as limiting factor. In this study we used analyte concentrations around 50 μM, which is 2 orders of magnitude lower than the minimum concentration required for typical potentiometric p*K*_a_ measurements. Theoretically, low analyte concentrations lead to more accurate p*K*_a_ measurements by minimizing the potential for the solute to influence solvent properties. In the extreme, if it were possible to measure the p*K*_a_ at the infinite dilution of the analyte that would be the best. But of course, in practice the minimum analyte concentration is limited by the detection strength of the UV signal. The higher the analyte concentration the more it affects the solvent properties such as ionic strength and dielectric constant. Also, the risk of analyte aggregation or precipitation increases with higher concentration.

In UV-metric measurements, both water and methanol (when used as cosolvent) stock solutions were ionic strength adjusted with 150 mM KCl, but acid and base solutions were not. This means that throughout pH titration ionic strength slighly fluctuates, but on average ionic strength of samples were staying around 150-180 mM. By using ISA solutions the effect of salt concentration change on p*K*_a_ measurements was minimized.

If an analyte is soluble enough, UV-metric p*K*_a_ measurements in water should be preferred over cosolvent methods, since p*K*_a_ measurement in water is more direct. For p*K*_a_ determination via cosolvent extrapolation using methanol, the apparent p*K*_a_s (p_s_K_a_) in at least three different methanol:water ratios must be measured, and the p*K*_a_ in 0% cosolvent computed by Yasuda-Shedlovsky extrapolation. The number and spread of p_s_K_a_ measurements and error in linear fit extrapolation influences the accuracy of p*K*_a_s determined by this approach. To test that UV-metric methods with or without cosolvent have indistinguishable performance, we collected p*K*_a_ values for 17 SAMPL6 compounds and pyridoxine with both methods. Figure 7 shows there is good correlation between both methods and the mean absolute deviation between two methods is 0.12 (95% CI [0.07, 0.18]). The mean deviation between the two sets is ‐0.04 (95% CI [-0.12, 0.03]), showing there is no significant bias in cosolvent measurements as the 95% CI includes zero. The largest absolute deviation observed was 0.41 for SM06.

### Impact of impurities to UV-metric p*K*_**a**_ measurements

Precisely how much the presence of small amounts of impurities impact UV-metric p*K*_a_ measurements is unpredictable. For an impurity to alter UV-metric p*K*_a_ measurements, it must possess a UV-chromophore and a titratable group in the vicinity of the chromophore—otherwise, it would not interfere with absorbance signal of the analyte. If a titratable impurity *does* possess a UV-chromophore, UV multiwavelength absorbance from the analyte and impurity will be convoluted. How much the presence of impurity will impact the multiwavelength absorbance spectra and p*K*_a_ determination depends on the strength of the impurity’s molar absorption coefficient and concentration, relative to the analyte’s. In the worst case scenario, an impurity with high concentration or strong UV absorbance can shift the measured p*K*_a_ value or create the appearance of an extra p*K*_a_. As a result, it is important to use analytes with high purities to obtain high accuracy p*K*_a_ measurements. Therefore, we confirmed the purities of SAMPL6 compounds with LC-MS.

### Interpretation of UV-metric p*K*_a_ measurements

Multiwavelength absorbance analysis on the Sirius T3 allows for good resolution of p*K*_a_s based on UV-absorbance change with respect to pH, but it is important to note that p*K*_a_ values determined from this method are often difficult to assign as either microscopic or macroscopic in nature. This method potentially produces *macroscopic* p*K*_a_s for polyprotic compounds. If multiple microscopic p*K*_a_s have close *pK*_*a*_ values and overlapping changes in UV absorbance spectra associated with protonation/deprotonation, the spectral analysis could produce a single macroscopic p*K*_a_ that represents an aggregation of multiple microscopic p*K*_a_s. An extreme example of such case is demonstrated in the simulated macrostate populations of cetirizine that would be observed with UV-metric titration (Figure 2).

If protonation state populations observed via UV-metric titrations (such as in Figure 3B) are composed of a single microstate, experimentally measured p*K*_a_s are indeed microscopic p*K*_a_s. Unfortunately, judging the composition of experimental populations is not possible by just using UV-metric or pH-metric titration. Molecules in the SAMPL6 p*K*_a_ challenge dataset with only one p*K*_a_ value measured in the 2–12 range could therefore be monoprotic (possessing a single titratable group that changes protonation state by gain or loss of a single proton over this pH range) or polyprotic (gaining or losing multiple protons from one or more sites with overlapping microscopic p*K*_a_ values). Similarly, titration curves of molecules with multiple experimental p*K*_a_s may show well-separated microscopic p*K*_a_s or macroscopic experimental p*K*_a_s that are really composites of microscopic p*K*_a_s with similar values. Therefore, without additional experimental evidence, UV-metric p*K*_a_s should not be assigned to individual titratable groups.

Sometimes it can be possible to assign p*K*_a_s to ionizable groups if they produce different UV-absorbance shifts upon ionization, but it is not a straight-forward analysis and it is not a part of the analysis pipeline of Sirius T3 Refine Software. Such an analysis would require fragmentation of the molecule and determining how UV-spectra of each chromophore changes upon ionization in isolation.

UV-metric p*K*_a_ values for nearly all compounds in our dataset with multiple resolvable p*K*_a_s are well-separated (more than 4 p*K*_a_ units), except for SM14 and SM18. Tam et al. states that spectrophotometric p*K*_a_ values of multiprotic molecules can be unambiguously assigned to the functional groups as microscopic p*K*_a_s "if the pKa values are at least 4 pH units apart (i.e. ***p**K***_*a*,2_** - ***p**K***_*a*,1_** ≥ 4)" based on general knowledge of functional groups and consideration of electronic and inductive effects [28]. In this study, we refrained from reporting such a knowledge-based assignment of p*K*_a_ values to functional groups without experimental evidence.

Determination of the exact microstates populated at different pH values via NMR can provide a complementary means of differentiating between microscopic and macroscopic p*K*_a_s in cases where there is ambiguity. As determination of protonation microstates via NMR is very laborious, we were only able to characterize microstates of two molecules: SM07 and SM14.

In UV-metric p*K*_a_ measurements with cosolvent, the slope of the Yasuda-Shedlovsky extrapolation can be interpreted to understand if the p*K*_a_ has dominantly acidic or basic character. As the methanol ratio is increased, p_s_*K*_a_ values of acids increase, while p_s_*K*_a_ values for bases decrease. However, it is important to remember that if the measured p*K*_a_ is macroscopic, acid/base assignment from cosolvent p_s_K_a_ trends is also a macroscopic property, and should not be used as a guide for assigning p*K*_a_ values to functional groups [60].

### NMR microstate characterization

The goal of NMR characterization was to collect information on microscopic states related to experimental p*K*_a_ measurements, i.e., determine exact sites of protonation. p*K*_a_ measurements performed with spectrophotometric method provide macroscopic p*K*_a_ values, but do not provide information on the specific site(s) of protonation. Conversely, most computational prediction methods primarily predict microscopic p*K*_a_ values. Protonation sites can be determined by NMR methods, although these measurements are very laborious in terms of data collection and interpretation compared to p*K*_a_ measurements with the automated Sirius T3. Moreover, not all SAMPL6 molecules were suitable for NMR measurements due to the high sample concentration requirements (for methods other than proton NMR, such as ^13^C and ^15^N based 2D experiments) and limiting analyte solubility. Heavy atom spectra that rely on natural isotope abundance require high sample concentrations (preferably in the order of 100 mM). It is possible that drug or drug-fragment-like compounds, such as the compounds used in this study, have insufficient aqueous solubility, limiting the choice of solvent and pH. It may be necessary to use organic cosolvents to prepare these high concentration solutions or only prepare samples at pH values that correspond to high solubility states (e.g., when the charged state of the compound is populated).

We performed NMR based microstate characterization only for SM07 and SM14. We were able to identify the order of dominant protonation microstates, as shown in Figure 8. These pairs of microstates and the order of microscopic transitions can be associated with experimental p*K*_a_s determined by UV-metric titrations, under the assumption that different organic solvents used in NMR measurements will have negligible effect on the sequence of microstates observed as the medium was titrated with acid, although shift in p*K*_a_ values is expected. NMR measurements for SM07 and SM14 were done in water:methanol (1:2 (v/v)) and acetonitrile solutions, respectively. On the other hand, p*K*_a_ values of these two compounds were determined by UV-metric titrations in ISA water. Additional UV-metric p*K*_a_ measurements of these compounds with methanol as a cosolvent showed that their p_s_K_a_ values decreased as the cosolvent ratio increased (i.e., dielectric constant decreased) as expected from base titration sites. Identification of SM07 and SM14 titratable sites type as base is consistent between NMR based models and UV-metric cosolvent titrations. The order of microstates observed in the titration of NMR samples are very likely to corresponds to the dominant microstates associated with UV-metric p*K*_a_ measurements. N-12 of SM07 was observed as the only protonation site of SM07 during TFA-*d* titration up to 5 equivalents which supports that SM07 is mono-protic and UV-metric p*K*_a_ value 6.08 ± 0.01 corresponds to microscopic protonation of N-12. For SM14, two protonation sites were observed (N-16 and N-9, in the order of increasing p_s_K_a_). Microstate pairs shown in Figure 8B were determined as dominant contributors to UV-metric p*K*_a_s 2.58±0.01 and 5.30±0.01, although minor microspecies with very low populations (undetected in NMR experiments) could be contributing to the macroscopic p*K*_a_ values observed by the UV-metric method.

In addition to SM07, there were five other 4-aminoquinazoline derivatives in the SAMPL6 set: SM02, SM04. SM09, SM12, and SM13. For these series, all the potential titratable sites are located in 4-aminoquinazoline scaffold and there are no other additional titratable sites present in these compounds compared to SM07. Therefore, based on structural similarity, it is reasonable to predict that N-12 is the microscopic protonation site for all of these compounds. We can infer that UV-metric p*K*_a_ values measured for the4-aminoquinazoline series are also microscopic p*K*_a_s and they are related to the protonation of the same quinazoline nitrogen with the same neutral background protonation states as shown for SM07 in Figure 8A.

### Recommendations for future p*K*_a_ prediction challenges

Most high-throughput p*K*_a_ measurement methods measure macroscopic p*K*_a_s. One way to circumvent this problem is to confine our interest in future p*K*_a_ challenges to experimental datasets containing only monoprotic compounds if UV-metric or pH-metric p*K*_a_ measurements are the method of choice, allowing unambiguous assignment of p*K*_a_ values to underlying protonation states. However, it is important to consider that multiprotic compounds are common in pharmaceutically interesting molecules, necessitating the ability to model them reliably. It might also be interesting to select a series of a polyprotic compounds and their monoprotic fragments, to see if they can be used to disambiguate the p*K*_a_ values.

Although relatively efficient, UV-metric p*K*_a_ measurements with the Sirius T3 do not provide structural information about microstates. Even the acid-base assignment based on direction of p_s_K_a_ shift with cosolvent is not a reliable indicator for assigning experimental p*K*_a_ values to individual functional groups in multiprotic compounds. On the other hand, most computational p*K*_a_ prediction methods output microscopic p*K*_a_s. It is therefore difficult to use experimental macroscopic p*K*_a_ values to assess and train microscopic p*K*_a_ prediction methods directly without further means of annotating macroscopic-microscopic correspondence. It is not straight-forward to infer the underlying microscopic p*K*_a_ values from macroscopic measurements of a polyprotic compound without complementary experiments that can provide structural information. Therefore, for future data collection efforts for evaluation of p*K*_a_ predictions, if measurement of p*K*_a_s via NMR is not possible, we advise supplementing UV-metric measurements with NMR characterization of microstates to show if observed p*K*_a_s are microscopic (related to a single group) or macroscopic (related to dissociation of multiple groups), as performed for SM07 and SM14 in this study.

Another source of complexity in interpreting macroscopic p*K*_a_ values is how the composition of macroscopic p*K*_a_s can change between different experimental methods as illustrated in Figure 2. Different subsets of microstates can become indistinguishable based on the type of signal the experimental method is constructed on. In potentiometric titrations, microstates with the same total charge are indistinguishable and are observed as one macroscopic population. In spectrophotometric p*K*_a_ measurements, the factor that determine if microstates can be resolved is not charge. Instead, microstates whose populations, and therefore UV-absorbance spectra, change around the same pH value become indistinguishable.

The "macroscopic" label is commonly ascribed to transitions between different ionization states of a molecule (all microstates that have the same total charge form one macrostate), but this definition only applies to potentiometric methods. In UV-absorbance based methods, the principle that determines which microstates will be distinguishable is not charge or number of bound protons, but molecular absorbance changes, and how closely underlying microscopic p*K*_a_ values overlap. To compare experimental macroscopic p*K*_a_ and microscopic computational predictions on common ground, the best solution is to compute "predicted" macroscopic p*K*_a_ values from microscopic p*K*_a_s based on the detection limitations of the experiment. A disadvantage of this approach is that experimental data cannot provide direct guidance on microscopic p*K*_a_ resolution for improving p*K*_a_ prediction methods.

Since analyte purity is critical for accuracy, necessary quality control experiments must be performed to ensure at least 90% purity for UV-metric p*K*_a_ measurements. Higher purities may be necessary for other methods. For potentiometric methods, knowing the stoichiometry of any counterions present in the original powder stocks is also necessary. Identity of counterions also needs to be known to incorporate titratable counterions, e.g. ammonia in the titration model.

For the set of SAMPL6 p*K*_a_ challenge compounds, we could not use potentiometric p*K*_a_ measurements due to the low aqueous solubility of many of these compounds. The lowest solubility observed *somewhere* in the experimental pH range of titration is the limiting factor, since for accurate measurements the analyte must stay in the solution phase throughout the entire titration. Since the titration pH range is determined with the goal of capturing all ionization states, the analyte is inevitably exposed to pH values that correspond to low solubility. Neutral and zwitterionic species can be orders of magnitude less soluble than ionic species. If a compound has a significantly insoluble ionization state, the pH range of titration could be narrowed to avoid precipitation, but it would limit the range of p*K*_a_ values that could be accurately measured.

For future p*K*_a_ challenges with multiprotic compounds, if sufficient time and effort can be spared, it would be ideal to construct an experimental p*K*_a_ dataset using experimental methods that can measure microscopic p*K*_a_s directly, such as NMR. In the present study, we were only able to perform follow up NMR microstate characterization of two compounds because we relied on intrinsically low-sensitivity and time-consuming ^1^H-^15^N HMBC experiment at natural abundance of ^15^N nuclei. ^1^H-^15^N HMBC experiments of SM07 and SM14 required high analyte concentrations and thus the use of organic solvents for solubility. Alternatively, it might be possible to determine microstates with ^1^H-NMR by analyzing chemical shift changes of reporter protons [21] in aqueous solutions with lower analyte concentrations and with much higher throughput than ^15^N-based experiments. However, it should be noted that ^1^H NMR titration data may not always be sufficient for unambiguous microstate characterization. In this case, other reporter nuclei such as ^13^C, ^19^F and ^31^P can be used where appropriate to supplement ^1^H data To prepare sample solutions for NMR at specific pH conditions, the Sirius T3 can be used to automate the pH adjustment of samples. Another advantage of using the Sirius T3 for NMR sample preparation includes preparing ionic strength adjusted NMR samples and minimizing consumption of the analyte since small volumes (as low as 1.5 mL) of pH adjusted solutions can be prepared.

In the future p*K*_a_ challenges, it would be especially interesting to expand this exercise to larger and more flexible drug-like molecules. p*K*_a_ values are environment dependent and it would be useful to be able to predict p*K*_a_ shifts based on on ionic strength, temperature, lipophilic content, with cosolvents or in organic solvents. Measuring the p*K*_a_ of molecules in organic solvents would be useful for guiding process chemistry. To test such predictions, special p*K*_a_ experiments would need to be designed to measure p*K*_a_s under different conditions.

The next iteration of the SAMPL log *D* prediction challenge will include a subset of compounds from p*K*_a_ challenge. We therefore envision that the collected dataset of p*K*_a_ measurements will also be of use for this challenge. Experimental p*K*_a_ values will be provided as an input to separate the p*K*_a_ prediction issue from other problems related to log *D* predictions. We expect that the experimental p*K*_a_s can be used as an indication if protonation states need to be taken into account for a log *D* prediction at a certain pH and for the validation of protonation state population predictions in the aqueous phase. Even for compounds for which microstates were not experimentally determined, macroscopic p*K*_a_ value can serve as an indicator of how likely it is that protonation states will have a significant effect on the log *D* of a molecule. Additionally, the information from NMR experiments in this study provided the site of protonation for six 4-aminoquinazoline compounds, which could be incorporated as microstate information for log *D* predictions. For predicting log *D* we suggest as a rule of thumb to include protonation state effects for p*K*_a_ values at least within 2 units of the pH of the log *D* experiment. p*K*_a_ values of six 4-aminoquinazoline compounds in this study were determined to be within 2 p*K*_a_ units from 7.

### Conclusion

This study reports the collection of experimental data for the SAMPL6 p*K*_a_ prediction challenge. Collection of experimental p*K*_a_ data was performed with the goal of evaluating computational p*K*_a_ predictions, therefore necessary quality control and uncertainty propagation measures were incorporated. The challenge was constructed for a set of fragment-like and drug-like small molecules, selected from kinase-targeted chemical libraries, resulting in a set of compounds containing heterocycles frequently found in FDA-approved kinase inhibitors. We collected p*K*_a_ values for 24 compounds with the Sirius T3 UV-metric titration method, which were then used as the experimental reference dataset for the SAMPL6 p*K*_a_ challenge. For compounds with poor aqueous solubilities we were able to use the Yasuda-Shedlovsky extrapolation method to measure p*K*_a_ values in the presence of methanol, and extrapolate to a purely aqueous phase.

In our work, we highlighted the distinction between microscopic and macroscopic p*K*_a_s which is based on the experimental method used, especially how underlying microstate composition can be different for macroscopic p*K*_a_ values measured with UV-metric vs pH-metric titration methods. We discuss how macroscopic p*K*_a_ values, determined by UV, introduce an identifiability problem when comparing to microscopic computational predictions. For two compounds (SM07 and SM14) we were able to alleviate this problem by determining the sequence of microscopic protonation states using ^1^H-^15^N HMBC experiments. Microstates of five other compounds with 4-aminoquinazoline scaffold were inferred based on the NMR characterization of SM07 microstates which showed that it is monoprotic.

The collected experimental data constitute a potentially useful dataset for future evaluation of small molecule p*K*_a_ predictions, even outside of SAMPL challenges. We expect that this data will also be useful for participants in the next SAMPL challenge on small molecule lipophilicity predictions.

## Code and data availability

- SAMPL6 p*K*_a_ challenge instructions, submissions, experimental data and analysis is available at https://github.com/MobleyLab/SAMPL6
- Python scripts used for compound selection are available at **compound_selection** directory of https://github.com/choderalab/sampl6-physicochemical-properties

## Overview of supplementary information

Supplementary tables and figures appearing in SI document:

- TABLE SI 1: Procurement details of SAMPL6 compounds
- TABLE SI 2: Selection details of SAMPL6 compounds
- TABLE SI 3: p*K*_a_ results of replicate experiments CSV
- TABLE SI 4: p*K*_a_ results of water and cosolvent replicate experiments CSV
- TABLE SI 5: p*K*_a_ mean and SEM results of water and cosolvent replicate experiments
- TABLE SI 6: Summary of LC-MS purity results
- FIGURE SI 1 - 24: LC-MS Figures
- FIGURE SI 25-35: NMR characterization of SM07 microstates
- FIGURE SI 36-54: NMR characterization of SM14 microstates

Additional files:

- Sirius T3 reports for all measurements: `supplementary_files.zip`

### Author Contributions

Conceptualization, MI, JDC, TR, ASR, DLM ; Methodology, MI, DL, IEN ; Software, MI, ASR ; Formal Analysis, MI ; Investigation, MI, DL, IEN, HW, XW, MR; Resources, TR, DL; Data Curation, MI ; Writing-Original Draft, MI, JDC, IEN; Writing - Review and Editing, MI, DL, ASR, IEN, HW, XW, MR, GEM, DLM, TR, JDC; Visualization, MI, IEN ; Supervision, JDC, TR, DLM, GEM, AAM ; Project Administration, MI ; Funding Acquisition, JDC, DLM, TR, MI.

## Acknowledgments

MI, ASR, and JDC acknowledge support from the Sloan Kettering Institute. JDC acknowledges support from NIH grant P30 CA008748. MI, JDC, ASR, and DLM gratefully acknowledge support from NIH grant R01GM124270 supporting SAMPL blind challenges. MI acknowledges support from a Doris J. Hutchinson Fellowship. DLM appreciates financial support from the National Institutes of Health (1R01GM108889-01), the National Science Foundation (CHE 1352608). IEN acknowledges support from the MRL Postdoctoral Research Program. The authors are extremely grateful for the assistance and support from the MRL Preformulations and NMR Structure Elucidation groups for materials, expertise, and instrument time, without which this SAMPL challenge would not have been possible. MI and DL are grateful to Pion/Sirius Analytical for their technical support in the planning and execution of this study. We are especially thankful to Karl Box (Sirius Analytical) for the guidance on optimization and interpretation of p*K*_a_ measurements with the Sirius T3, as well as feedback on the manuscript. We thank Brad Sherborne (MRL; ORCID: 0000-0002-0037-3427) for his valuable insights at the conception of the p*K*_a_ challenge and connecting us with TR and DL who were able to provide resources for experimental measurements. We acknowledge Paul Czodrowski (Merck KGaA; ORCID: 0000-0002-7390-8795) who provided feedback on multiple stages of this work: challenge construction, purchasable compound selection, and manuscript. We acknowledge contributions from Caitlin Bannan who provided feedback on experimental data collection and structure of p*K*_a_ challenge from a computational chemist’s perspective. We are also grateful to Marilyn Gunner (CCNY) for her feedback on this manuscript. We thank anonymous reviewers for their input and constructive comments that improved this manuscript. MI, ASR, and JDC are grateful to OpenEye Scientific for providing a free academic software license for use in this work. The content is solely the responsibility of the authors and does not necessarily represent the official views of the National Institutes of Health.

